# Disrupted visual cortical plasticity in early neurodegeneration

**DOI:** 10.1101/2020.11.02.365767

**Authors:** Amalia Papanikolaou, Fabio R. Rodrigues, Joanna Holeniewska, Keith Phillips, Aman B. Saleem, Samuel G. Solomon

## Abstract

Neurodegeneration is a hallmark of many dementias and is thought to underlie a progressive impairment of neural plasticity. How neurodegeneration affects plasticity in neural circuits is not known. We therefore characterised the impact of tau-driven neurodegeneration on plasticity in the visual system, where normal function is well understood. We studied a very simple form of visual plasticity that allowed us to track both long timescales (across days) and shorter timescales (over minutes). We recorded the local field potential in the primary visual cortex of rTg4510 transgenic mice, a mouse model of tauopathy, while animals were repeatedly exposed to the same stimulus over the course of 9 days. We studied animals at early stages of neurodegeneration (5 months old) and at a more advanced stage where pathology is evident (8 months). We found that both short- and long-term visual plasticity were already disrupted at early stages of neurodegeneration, and were further reduced in older animals, such that it was abolished in mice expressing the mutant tau. Additionally, we found that visually evoked behaviours were disrupted in both younger and older mice expressing the mutant tau. Our results show that visual cortical plasticity and visually evoked behaviours are disrupted in the rTg4510 model of tauopathy, even at early stages of neurodegeneration. This simple measure of neural plasticity may help understand how neurodegeneration disrupts neural circuits, and offers a translatable platform for detection and tracking of the disease.

**Highlights:**

1. *<u>Visual plasticity is disrupted at early stages of neurodegeneration in rTg4510 mice</u>*
2. *<u>Visual plasticity is reduced in older animals, particularly during neurodegeneration</u>*
3. *<u>Instinctive visual behaviours are reduced in neurodegeneration</u>*
4. *<u>Short-term visual plasticity is reduced in neurodegeneration</u>*

## Introduction

Neurodegenerative diseases are known to affect neural plasticity and memory. While the cellular and molecular basis of many neurodegenerative diseases is increasingly clear, how these diseases alter the plasticity of neural circuits is not well understood. Poor understanding of neurodegeneration’s impact on circuit plasticity may reflect the fact that previous research has concentrated on plasticity in circuits thought important for memory and cognition, where even normal function is not well understood. But plasticity and neurodegeneration are a feature of all neural circuits, even those involved in simpler processes. For example, in addition to memory deficits, visual deficits are also common in dementia, and are particularly acute in patients with posterior cortical atrophy (PCA), which affects many sufferers of Alzheimer’s disease (AD) (for a review see (Crutch et al., 2016)). The neural bases of these effects in PCA patients is thought to be tau-related pathological changes in posterior cortices such as the occipital lobe (Hof et al., 1990; Ossenkoppele et al., 2015; Tang-Wai et al., 2004). Similarly, most animal models of neurodegenerative diseases exhibit widespread pathological impact, including sensory cortical areas such as the visual cortex.
We characterised the relationship between neurodegeneration and visual plasticity in a commonly used and well-characterised transgenic mouse model of neurodegeneration, the rTg4510 model of tauopathy. rTg4510 mice overexpress a mutant form of the human microtubule-associated protein tau (Ramsden et al., 2005; Santacruz et al., 2005). While in the healthy brain tau plays a role in the stabilization and assembly of microtubules, primarily in axons, in tauopathy it is hyperphosphorylated and forms abnormal aggregates in the soma and dendrites called neurofibrillary tangles (NFT) (Grundke-Iqbal et al., 1986; Mondragon-Rodriguez et al., 2008). These aggregates may disrupt internal neuronal processes, leading to neuronal degeneration and cell death. The rTg4510 mice develop progressive NFT, neuronal loss and concomitant cognitive deficits (Ramsden et al., 2005; Yue et al., 2011). High levels of mutant tau emerge in the hippocampus and neocortex between 2 and 4 months of age. NFT are present by 4.5 months in hippocampus and 7-8 months in neocortex (Ramsden et al., 2005; Santacruz et al., 2005). Cortical cell loss occurs slightly later, at about 8.5 months (Spires et al., 2006). The rTg4510 model therefore allows the characterisation of functional plasticity across different stages along the progression of tauopathy.

We studied a very simple form of visual plasticity, by measuring the visual cortical response to repeated presentation of a flickering pattern. Over several minutes, repeated presentation of a pattern usually suppresses response to that pattern, a classical effect of sensory adaptation (Solomon and Kohn, 2014; Webster, 2015). Over several days, however, repeated presentation of a visual stimulus can instead increase response, involving a sleep-dependent process called stimulus response potentiation, or SRP (Aton et al., 2014; Cooke et al., 2015; Frenkel et al., 2006). SRP is a form of longterm plasticity that resembles canonical long-term synaptic potentiation (LTP), with which it shares mechanisms (Cooke et al., 2015). Visual cortical plasticity may therefore offer a simple and direct measurement of experience-dependent plasticity in brain circuits, over multiple relevant timescales, that could help link functional and structural changes during neurodegeneration.

We used this simple paradigm to understand how brain plasticity is altered in the rTg4510 model of tauopathy. We studied animals at early stages of neurodegeneration (~5 months old) and at a more advanced stage where clear degeneration has taken place in the cortex (~8 months). Mice expressing the mutant Tau were compared with littermates that were fed doxycycline to suppress the expression of the transgene (Blackmore et al., 2017; Santacruz et al., 2005; Spires et al., 2006). We found that both short- and long-term visual plasticity are reduced at early stages of neurodegeneration. Visual plasticity was further reduced in older animals, such that it was abolished at later stages of neurodegeneration. Additionally, we found that mice expressing the mutant tau showed reduced visually evoked behaviours. Our results show that visual cortical plasticity and visually evoked behaviours are disrupted in the rTg4510 model of tauopathy, even at early stages of neurodegeneration.

## Results

We measured the local field potential (LFP) from the primary visual cortex (V1) of 50 rTg4510 mice, using electrodes targeted to layer 4. Half of the mice were placed on a doxycycline diet from 2 months of age, to suppress the expression of mutant tau (referred as Tau−). We used different animals to study early stages of neurodegeneration (~5 months) and more advanced stages (~8 months) (Fig. 1A). Mice fed a normal diet, and therefore continuing to express the mutant Tau (referred as Tau+) showed increased tau pathology (F=23.61, p=10^−4^, two-way ANOVA, Fig. 1B, Fig. S1) and a significant reduction in the overall brain weight compared to Tau− animals (F=5.79, p=0.02, Fig. 1B).

**Figure 1.**
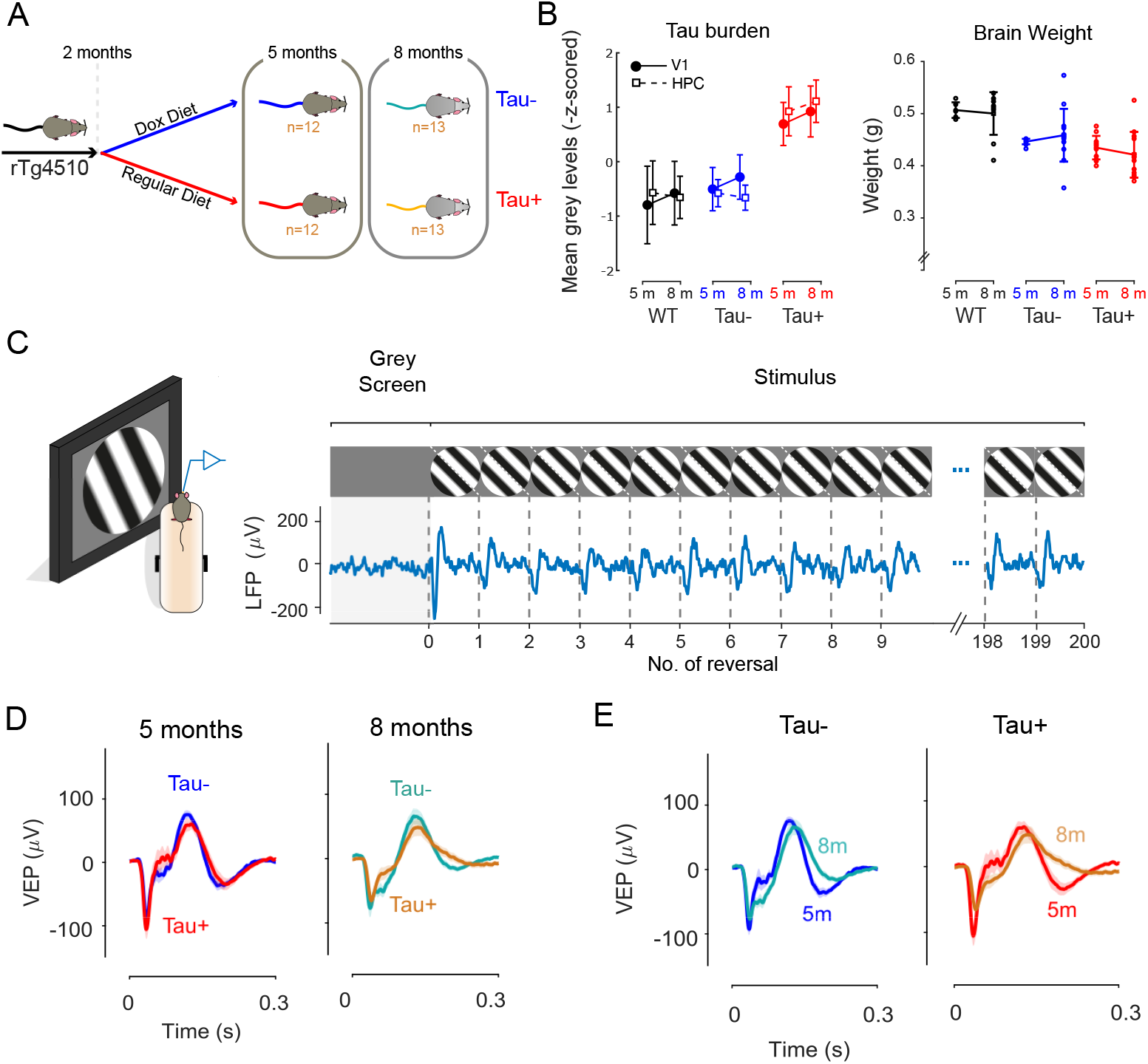
Visual evoked responses in the primary visual cortex are largely preserved in neurodegeneration. **A.** At 2 months of age, half the rTg4510 mice were treated with and fed doxycycline (Dox) to suppress the expression of the transgene and arrest further accumulation of Tau. **B.** Left, comparison of average tau burden for each group of animals calculated as the mean grey levels within selected regions of interest in V1 and hippocampus (HPC) (Methods). Data are represented as mean±2*SEM. Right, comparison of overall brain weights at 5 months (5 m) and 8 months (8 m) timepoints. Data are represented as mean±SEM. **C.** Mice were head-fixed and allowed to run on a styrofoam wheel. After 5 min exposure to a homogenous grey screen, a full-screen, contrast reversing, sinusoidal grating pattern was presented to the left monocular visual field. The grating reversed every 0.5s, and the animal was exposed to 10 blocks of 200 continuous reversals, with 30s presentation of a grey screen between blocks. We recorded the local field potential (LFP) in the right primary visual cortex (V1). The LFP trace shown is the average time course across 10 stimulus blocks on one day in one Tau− mouse. Each reversal generated a visual evoked potential (VEP), characterized by an initial negative deflection and subsequent positive one. **D.** Average VEP responses on the first day of recording for Tau− and Tau+ animals at 5 months (left) and 8 months (right). Shaded area represents the SEM. There was no significant difference in the size and shape of the VEPs between groups at either age (p=0.95, twoway ANOVA). **E.** Average VEP responses for Tau− (left) and Tau+ (right) mice obtained at 5 months and 8 months of age. VEPs were slightly reduced, and more sustained, for both Tau− and Tau+ mice at 8 months.

### Visual cortical responses in rTg4510 mice

We recorded the visual evoked potential (VEP) in response to a large sinusoidal grating, presented to the monocular visual field by a computer monitor. Mice were head-fixed but free to run on a styrofoam wheel. The contrast of the grating was flipped (reversed) every 0.5s, and the animals were exposed to 10 blocks of 200 reversals, with 30s of a grey screen presented between blocks. Each reversal generated a VEP, with an initial negative deflection rapidly followed by a positive one (Fig. 1C). We found that the amplitude of the VEP was similar in Tau− and Tau+ animals, for both 5- and 8-month old animals (Fig. 1D). We calculated the VEP amplitude as the difference between the positive and negative peaks of the VEP signal and found that the average amplitude was not different between genotypes (F=0, p=0.95, two-way ANOVA) or age groups (F=3.17, p=0.08). A Tukey’s multiple comparison test showed no significant difference between Tau− and Tau+ VEP amplitudes at 5 months (Tau− 5m: 165±11, Tau+ 5m: 184±16, mean±SEM, p=0.85) or 8 months (Tau− 8m: 155±19, Tau+ 8m: 134±16, mean±SEM, p=0.79). At 8 months, the VEP signal reduced slightly for Tau+ mice compared to their 5-month old counterparts (Fig. 1E), but the effect did not reach significance (p=0.16, Tukey’s test). The VEP signal was more sustained in 8-month old animals for both Tau− and Tau+ animals. Our results therefore show that basic visual cortical responses are largely preserved in the rTg4510 mouse model, even during advanced neurodegeneration.

### Visual plasticity is disrupted in mice expressing mutant Tau

We hypothesised that plasticity is more likely to be affected than basic visual response at early stages of neurodegeneration. Stimulus-Response Potentiation (SRP) is a form of cortical plasticity that can be measured in the VEP of mouse V1. SRP is induced by repeated exposure to a visual stimulus over several days (Frenkel et al., 2006). We used this simple paradigm to assess the impact of tauopathy on cortical plasticity. We obtained VEP measurements from 24 5-month old rTg4510 mice (12 Tau− and 12 Tau+) while they were exposed to a grating of one orientation (either 45° or −45° from vertical) over 9 days (thus becoming a familiar stimulus). On the first and last day of recordings (day 1 and day 9), five blocks of this familiar stimulus orientation were interleaved with five blocks of the orthogonal orientation (unfamiliar stimulus). On days 2-8, 10 blocks of the familiar stimulus were presented (Fig. 2A).

**Figure 2.**
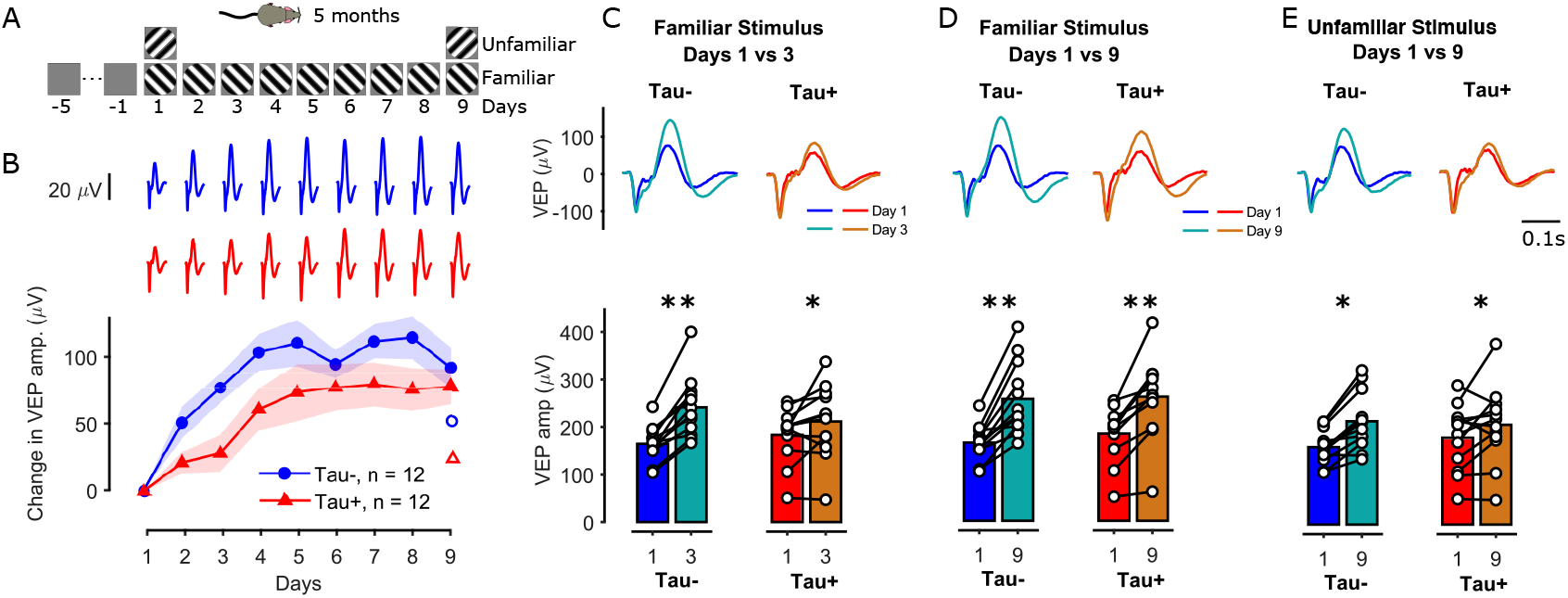
Visual plasticity is reduced at early stages of tauopathy. **A.** Mice (5 months old) were exposed to a grating of one orientation (either 45° or −45°; ‘familiar’ stimulus) for 9 days. On the first and last day of recordings, five blocks of this familiar stimulus were interleaved with five blocks of a grating (‘unfamiliar’) whose orientation was orthogonal to the familiar grating. On days 2-8, 10 blocks of the familiar stimulus were presented. **B.** Change in the VEP amplitude from day 1 for the familiar stimulus for Tau− (blue) and Tau+ (red) animals over the course of days. Tau+ mice showed a slower potentiation of the LFP signal compared to Tau− animals. The open symbols on the right show the change in VEP amplitude in response to the unfamiliar stimulus on day 9. **C-E.** Comparison of average VEPs (top) and VEP amplitudes (bottom) between: days 1 vs 3 for the familiar stimulus (**C**), days 1 vs 9 for the familiar stimulus (**D**), and days 1 vs 9 for the unfamiliar stimulus (**E**).

We found strong potentiation of the VEP signal in Tau− animals, and reduced and slower potentiation in Tau+ animals (Fig. 2). The VEP amplitude for Tau− mice reached a plateau around day 3-4 (Fig. 2B) while Tau+ mice showed a slower increase in the VEP amplitude, reaching a plateau around day 5-6. For example, the VEP amplitude of Tau− mice was 77±11 μV larger on day 3 than it was on day 1 (Fig. 2B,C; p=3.1*10^−6^, repeated measures ANOVA, Tukey’s pairwise comparison), while Tau+ animals showed only a moderate increase by day 3 (28±13 μV, p=0.03). By day 9, VEP amplitude had increased substantially for both groups of mice (Fig. 2D, Tau−: 92±15 μV, p=1.5*10^−6^, Tau+: 78±13, p=1.6*10^−5^). Repeated measures ANOVA on days 1 and 3 revealed a significant day by phenotype interaction (F=7.75, p=0.01), while the same comparisons for days 1 and 9 showed a significant effect of day (F=72.1, p=2.1*10^−8^) but no interaction (F=0.5, p=0.48). The slower growth in VEP amplitude in Tau+ animals was mainly due to slower changes in the positive deflection of the VEP signal in Tau+ animals (Fig. S2b). The negative deflection of the VEP was slightly larger in Tau+ mice than in Tau− mice, and increased at a similar rate in both groups (Fig. S2c).

Repeated exposure to the familiar stimulus had less impact on VEP response to the unfamiliar stimulus (which was shown only on day 1 and day 9), for both groups (Fig. 2E, Tau−: unfamiliar VEP amp. change=54±10, p=0.2*10^−3^, Tau+: 15±27, p=0.03, Tukey’s test). In fact, growth in VEP responses to the unfamiliar stimulus on day 9 was comparable to growth in VEPs to the familiar stimulus on day 2 (Fig. 2B, Tau−: day 2 familiar VEP amp. change=51±12, p=0.89, Tau+: 21±8, p=0.72). Together these data suggest that the amplitude but not stimulus selectivity of response potentiation is affected in Tau+ animals.

Overall our results suggest that visual cortical plasticity is disrupted even at this early stage of neurodegeneration.

### Visual plasticity is reduced in older animals

We showed that visual plasticity is affected even at early stages of neurodegeneration in rTg4510 mice. We then asked whether functional deficits increase with age by examining visual cortical plasticity in 8-month old rTg4510 mice. We obtained LFP responses from 26 8-month old transgenic mice (13 Tau− and 13 Tau+) using the same visual paradigm described above (Fig. 3A).

**Figure 3.**
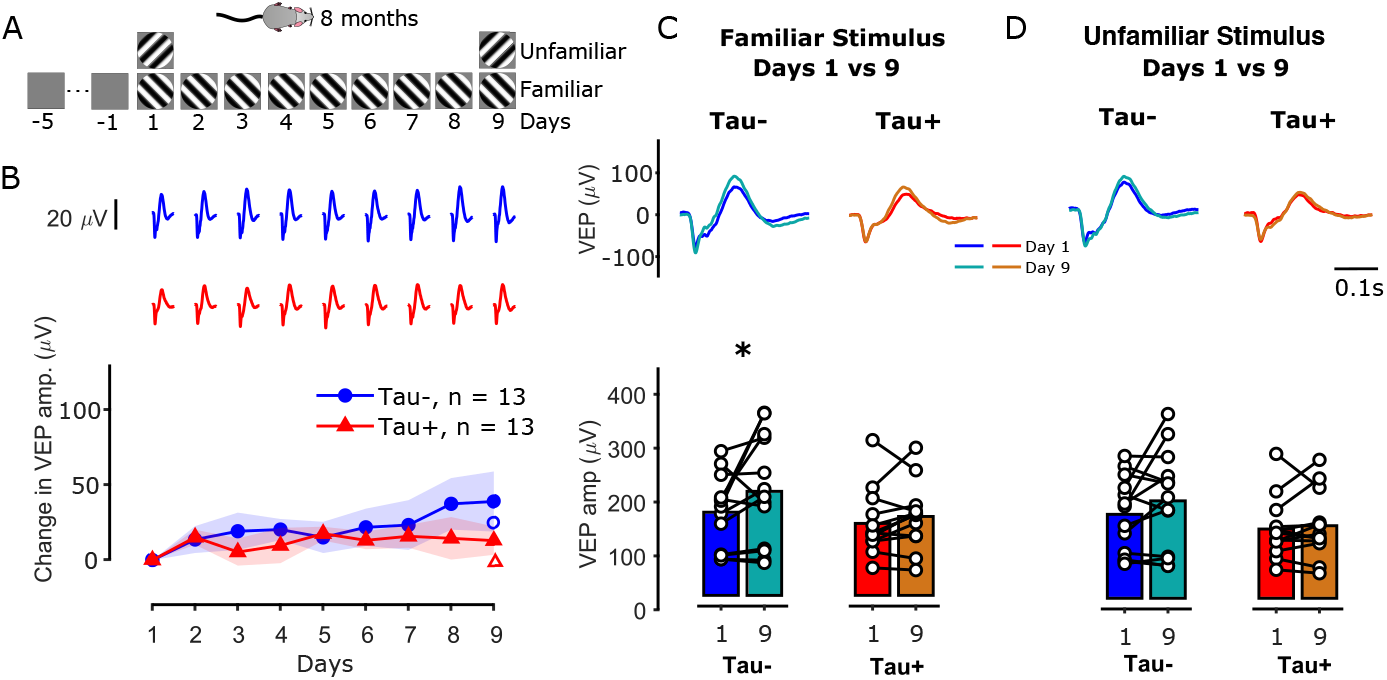
Visual plasticity diminishes with age. **A.** Mice (8 months old) were exposed to a grating of one orientation (either 45° or −45°; ‘familiar’ stimulus) for 9 days (same as in Fig. 2A). **B.** Difference in the VEP amplitude from day 1 for the familiar stimulus for Tau− (blue) and Tau+ (red) animals over the course of days. 8-month old Tau− mice showed a smaller potentiation of the LFP signal compared to 5-month old mice (Fig. 2B). Tau+ mice showed no significant potentiation. The open symbols on the right show the change in VEP amplitude in response to the unfamiliar stimulus on day 9. **C-D**. Comparison of average VEPs (top) and VEP amplitudes (bottom) between: days 1 vs 9 for the familiar stimulus (**C**), and days 1 vs 9 for the unfamiliar stimulus (**D**).

Visual plasticity was reduced in older animals (Fig. 3). By day 9, the VEPs of Tau− mice showed a small but significant increase relative to day 1 (Fig. 3C, VEP amp. change=39±20 μV, p=0.02), while Tau+ animals showed minimal change (13±9, p=0.41). No significant change was observed for the unfamiliar stimulus in either group (Fig. 3D, Tau−: VEP amp. change=25±16, p=0.72, Tau+: 6±9, p=0.63). Our results suggest that there may be a combined effect of mutant tau expression and age leading to a complete disruption of visual cortical plasticity.

### Similar response potentiation in Tau− and WT mice

Tau− mice were fed with a doxycycline diet to suppress mutant tau transgene expression from the age of 2 months. It is possible however that expression of mutant tau up to the age of 2 months, or a continued effect thereafter, could lead to functional deficits in these animals. For example, the reduced visual plasticity observed in 8-month old compared to 5-month old Tau− mice might be a pathological effect initiated by early tauopathy. We therefore compared the visual responses of Tau− mice with responses obtained from wild type (WT) littermates.

At 5 months, WT mice showed similar VEP potentiation to the Tau− mice (Fig. S3A). By day 3, WTs showed a significant increase in the VEP amplitude for the familiar stimulus relative to day 1 (VEP amp. change=57±11 μV, p=0.005), similar to Tau− mice (Fig. S3A). At 8 months, the VEP amplitude was on average larger in WT compared to Tau− mice (F=0.92, p=0.35, one-way ANOVA). As for Tau− mice, however, 8-month old WT mice showed reduced and slower potentiation of the LFP signal for the familiar stimulus compared to 5-month old mice (Fig. S3B). This suggests that the reduction in SRP at 8 months is largely an effect of age and not neurodegeneration in Tau− mice.

### Coincident changes in short-term visual plasticity

We have shown that long-term visual plasticity is disrupted in rTg4510 mice using a simple and well-established visual paradigm. Although SRP has been conventionally used as a measure of plasticity across days, it can also be used to measure changes within a day, or within a block. We therefore asked if there were also disruptions in plasticity at these shorter timescales. We analysed how responses changed within each day. We considered days 2-8 where only the familiar stimulus was presented (Fig. 4A).

**Figure 4:**
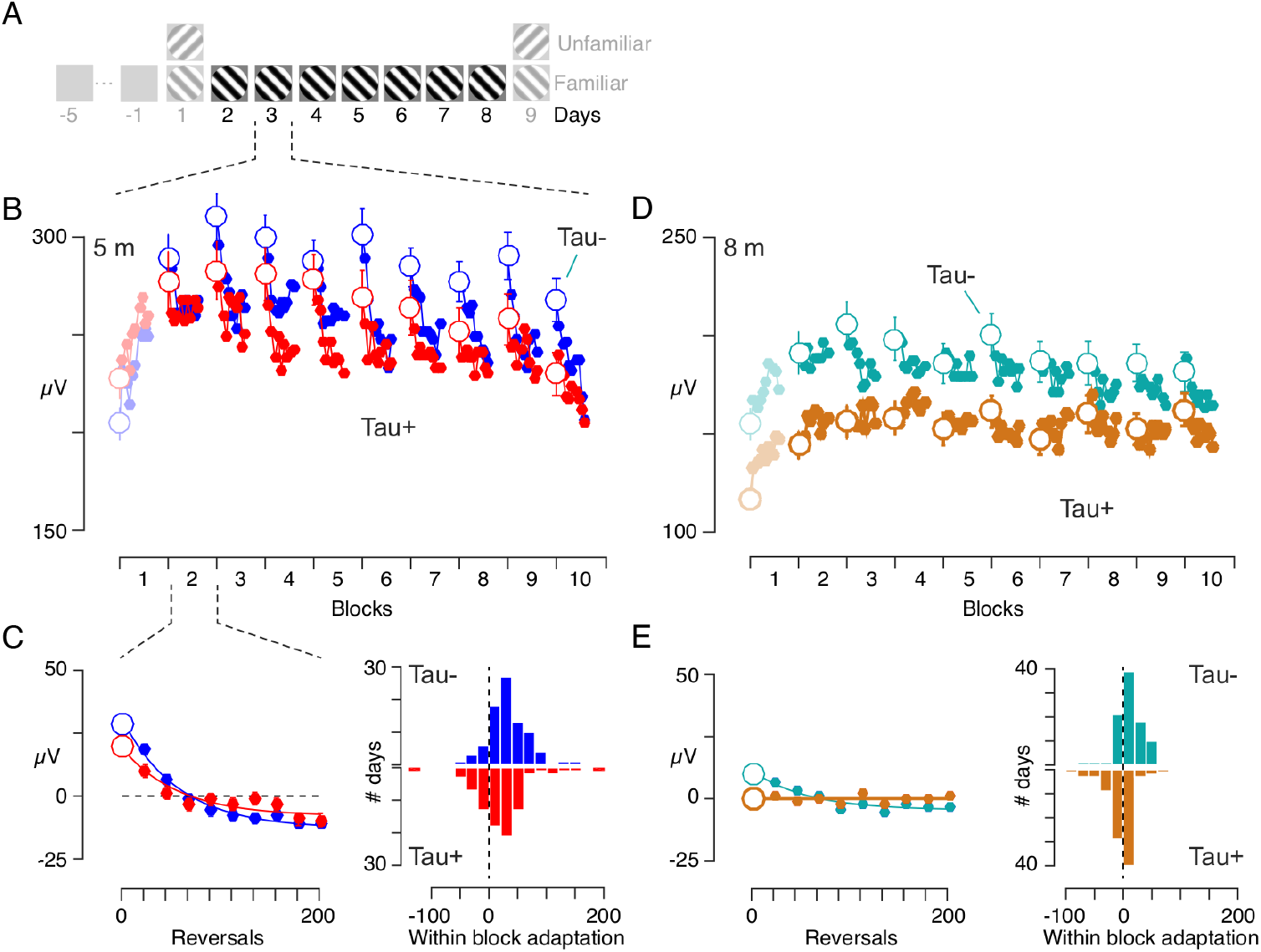
Short-term visual plasticity is reduced in rTg4510 mice expressing mutant tau. **A.** We measured intra-day effects on experimental days 2-8. On each day 10 blocks of the familiar stimulus were presented, each consisting of 200 phase reversals and separated by 30s of a homogenous grey screen. **B.** Average VEP amplitude as a function of block number for 5-month old Tau− (blue) and Tau+ (red) mice. The VEP amplitude was calculated from non-overlapping averages of 20 reversals. With the notable exception of the first block, VEPs showed a reduction of responses within each block, consistent with classic sensory adaptation effects. **C.** Left: Average VEP responses for blocks 2-10 on days 2-8, for Tau− and Tau+ animals. Right: To measure adaptation within a block, we subtracted the mean VEP amplitude from each block and then averaged across blocks 2-10. The within block effects (at a resolution of 20 reversals) were then fit with a decaying exponential function of fixed τ=7.6 reversals (see Methods), and the amplitude of the exponential was extracted. The histograms show the distribution of these fitted amplitude. At 5 months, T au+ animals showed reduced within-block adaptation compared to Tau− mice. **D-E.** Same as A,B for 8-month old animals. The within-block adaptation is reduced for Tau− mice at 8 months compared to 5 months. Tau+ mice showed no adaptation effect.

The amplitude of the VEPs reduced over the course of each block, consistent with classic sensory adaptation effects (Fig. 4B). We observed this suppressive adaptation effect in both Tau− and Tau+ animals, at least in blocks 2-10. The first block presented each day was instead characterised by an enhancement of responses, in both groups of animals. Averaging VEP responses across blocks 2-10 and days 2-8 showed smaller adaptation effects in Tau+ than Tau− mice (Fig. 4C). To quantify this, we fitted a decaying exponential function to the within-block time course of the VEP amplitude. We subtracted the mean VEP amplitude from each block and then averaged across blocks recorded on the same day, providing one estimate of adaptation’s effect for each day in each animal. At 5 months, Tau+ animals showed significantly reduced within-block adaptation compared to Tau− mice (Fig. 4C; Tau−: 34±32 μV, Tau+: 21±45, p=0.03, t-test). Adaptation effects were reduced in 8-month old Tau− mice compared to 5-month olds (Fig. 4D,E), and they were completely abolished in 8-month Tau+ mice (Fig. 4C,E; Tau−: 11±21, Tau+: −3±24, p=0.0001). Our results suggest that visual plasticity is disrupted in neurodegeneration even at time scales as short as several seconds.

Interestingly, unlike Tau− animals, 5-month and 8-month old WTs showed similar within block adaptation effects (Fig. S4, Fig. 4; 5m: 26±35 μV, 8m: 26±30, p=0.99, t-test), suggesting that while long term plasticity is reduced in older WT animals, short term plasticity is not. Reduced adaptation in older Tau− animals may reflect persistent impact of the initial tau accumulation before the onset of the doxycycline treatment (which was started at 2 months), subsequent incomplete suppression of the transgene, or age-dependent effects of other genetic disruptions in this mouse model (see Discussion).

### Differences in visual plasticity cannot be explained by differences in behavioural state

Animals were free to run on a styrofoam wheel during the recording session, and we observed epochs of running and variations in pupil size in all animals. Visual cortical responses, as well as responses earlier in the visual pathway, are known to vary with behavioural state as defined by running speed or pupil area (Niell and Stryker, 2010). Tau+ rTg4510 mice have also been reported to have abnormal locomotion behaviours (Blackmore et al., 2017; Cook et al., 2014; Wes et al., 2014). We therefore wanted to know if the differences observed in visual plasticity between Tau+ and Tau− animals might be explained by differences in behavioural state. We used two approaches to address this. We first refined our analyses by only considering VEP responses during either stationary (speed < 5cm/s) or running (speed > 5cm/s) epochs. We next used a model to evaluate the contribution of various parameters to the observed responses.

The VEP signal was smaller during running, in both Tau− and Tau+ mice (Fig. 5A). This reduction was seen across all days and animals (Fig. 5B), and was slightly more pronounced in Tau+ animals. To establish whether the animal’s arousal state was responsible for the differences observed in SRP between groups, we calculated the change in the VEP amplitude from day 1, but considering only stationary or only running epochs (Fig. 5C; Sup. Fig. S5B). The differences in SRP between Tau− and Tau+ mice were more pronounced when we considered only stationary epochs (Fig. 5C, S5C-D), so differences in running behaviour cannot explain the differences in SRP.

**Figure 5:**
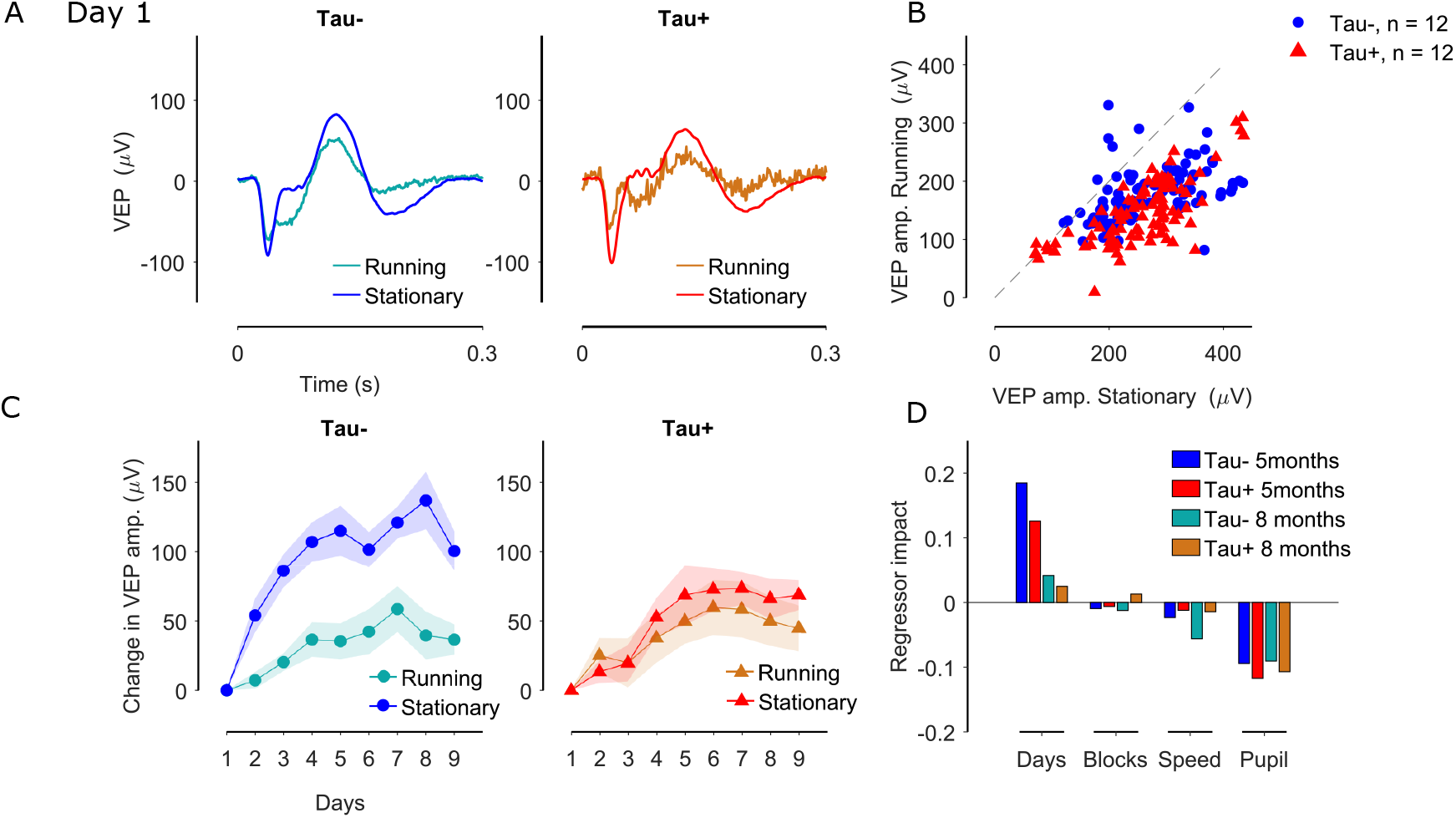
Differences in visual plasticity are not due to differences in behavioural state. **A.** VEP responses on day 1 for Tau− (blue) and Tau+ (red) 5-month old animals calculated during running (speed>5cm/s) and stationary epochs. Running reduces the VEP signal for both groups of mice. **B.** VEP amplitude during stationary versus running epochs for individual Tau− (blue circles) and Tau+ (red triangles) mice. Each symbol represents the VEP amplitude on one day in one animal. **C.** Difference in the VEP amplitude from day 1 for the familiar stimulus, for Tau− (left) and Tau+ (right) 5-month old animals, calculated for only stationary or only running epochs. VEP potentiation was evident in both groups of animals for both behavioural states. **D.** We fitted an elastic net regularization model to predict the VEP amplitude for each animal using days, block number, movement speed and pupil diameter as regressors. We assessed the impact of each regressor in predicting the VEP amplitude as the product between each learned weight and observed variable, divided by the predicted value across the entire regularisation path (of elastic net regression). The panel shows the average regressor impact for each genotype at 5 and 8 months of age.

Our analyses show that VEP amplitude is dependent on behavioural state, and varies within a day (Fig 4). These effects might combine to accentuate or mask plasticity across days. To assess the relative impact of time and behaviour on the VEP, we used an elastic net regularization model to predict the VEP amplitude for each animal using *Day*, *Block* number, movement *Speed* and *Pupil* diameter as regressors. We found *Day* to have a positive impact on the VEP amplitude, thus predicting the increase in VEP amplitude across days (Fig. 5D). Consistent with our observations on visual plasticity (Fig. 2,3), the impact of *Day* in predicting the VEP amplitude was greater for Tau− than Tau+ animals, for both 5-month old and 8-month old animals. Similarly, *Day* had a lower impact at 8 months than 5 months old. Block number had a negligible overall impact in predicting the VEP amplitude compared to the other predictors considered here. Increases in behaviour (defined by pupil diameter and speed) had a negative impact on the VEP amplitude, and this relationship was similar across phenotypes and age groups. In addition, there was no correlation between behaviour and days (Pupil: r = −0.03, Speed: r=0.07) suggesting that behaviour cannot explain the VEP increase across days. Therefore, our results suggest that the VEP potentiation to the familiar stimulus across days is an effect of experience to the visual stimulus and not an effect of changes in the behavioural state of the animals.

### Visual evoked behaviours are reduced in naive Tau+ mice

In animals, including mice, an unexpected or unfamiliar visual stimulus usually evokes instinctive behavioural responses. In head-fixed animals who are unable to run, these behavioural responses can include low amplitude muscle movements (‘fidgeting’). SRP in the mouse visual cortex has previously been associated with habituation of fidgeting (Cooke et al., 2015). To establish whether reduced plasticity in Tau+ mice is associated with reduced behavioural responses in our conditions, we assessed the impact of stimulus presentation on pupil diameter and movement speed, both at first exposure to the stimulus, and later when animals were more experienced.

Tau− (and WT) mice showed large dilation of the pupil and an increase in the movement speed at the onset of their very first exposure to the grating stimulus (day 1, block 1; Fig. 6A,B, Fig. S3). These responses quickly habituated, both within the session (Sup. Fig. S6) and over the course of days (Fig. 6C,D). By contrast, Tau+ mice showed no visual evoked behavioural responses to the onset of the stimulus (Fig. 6A,B). These group differences in visual evoked behaviours were clear at both 5 and 8 months of age.

**Figure 6:**
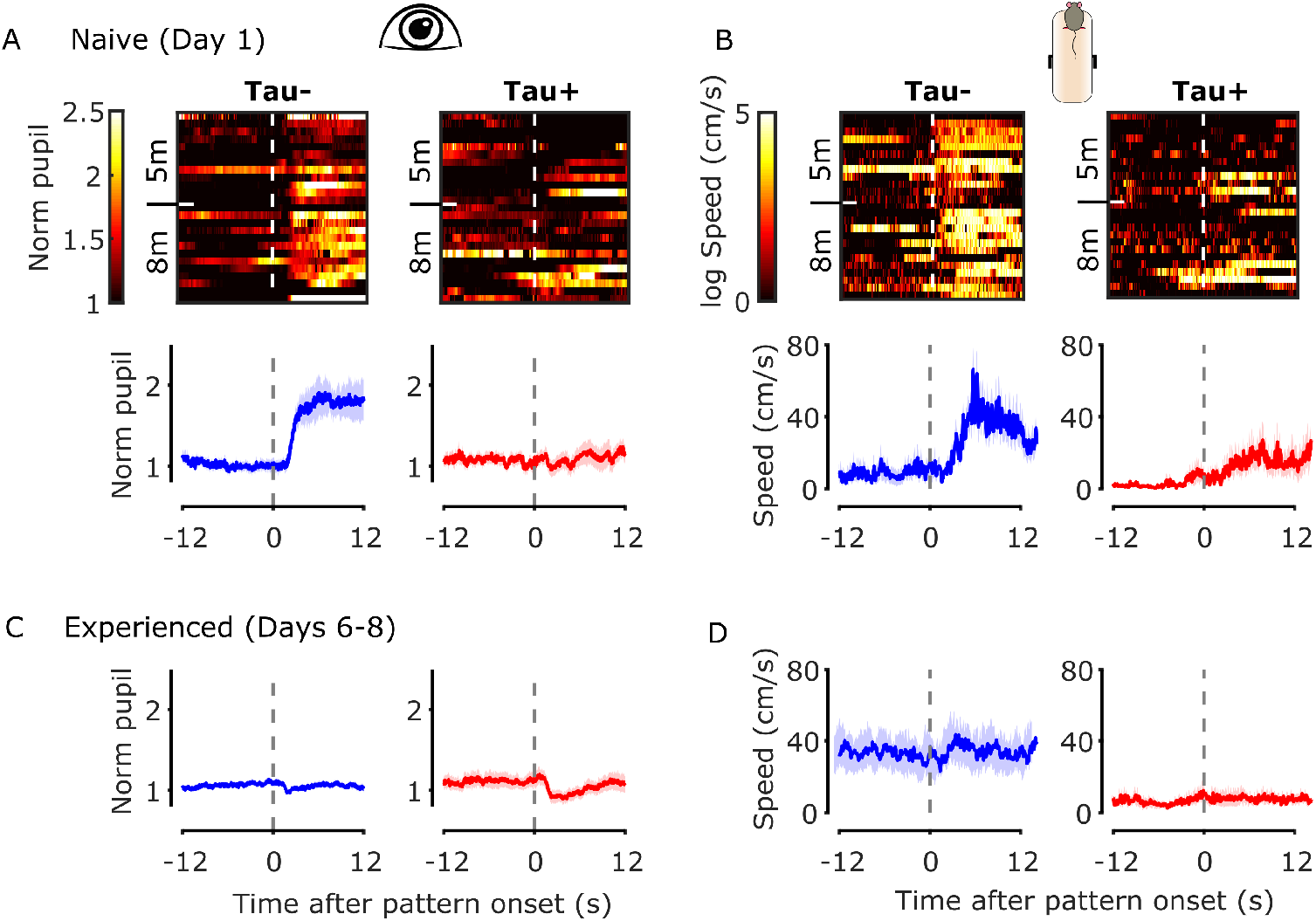
Visual evoked behaviours are reduced in Tau+ mice. **A.** *Top:* Images of the normalized pupil responses to the onset of the stimulus of the first block presented for naive animals (day 1; one animal per row). Pupil responses are normalized to the average pupil size during stationary epochs for 12s before the stimulus onset where mice were viewing a grey screen. *Bottom:* Mean (±SEM) pupil responses for naive Tau− (blue) and Tau+ (red) animals for both age groups. **B.** *Top:* Images of the natural logarithm of movement speed responses to the onset of the stimulus for naive animals (day 1). *Bottom:* Mean (±SEM) movement speed for naive Tau− (blue) and Tau+ (red) animals for both age groups. **C-D.** Average normalized pupil responses and average movement speed to the onset of the stimulus for experienced animals (days 6-8). Horizontal dashed lines indicate the stimulus onset.

Experienced Tau− animals (days 6-8) showed a small constriction of the pupil in response to the stimulus onset. This constriction was more pronounced in Tau+ animals (Fig. 6C). There was no change in the movement speed for both groups in response to the stimulus, although experienced Tau− mice showed larger average speed compared with the Tau+ mice (Fig. 6D).

Overall, our results suggest that visual evoked responses are disrupted even in early stages of neurodegeneration.

## Discussion

In this study, we evaluated visual cortical plasticity in the rTg4510 mouse model of neurodegeneration. We measured both short term suppression and longterm potentiation of the visual evoked LFP response in V1, in mice with mutant Tau expression (Tau+), or with that expression suppressed (Tau−). We made these measurements at two time points, at an early (5 months), and a more advanced (8 months) stage of neurodegeneration (Helboe et al., 2017; Ramsden et al., 2005; Santacruz et al., 2005; Spires et al., 2006), which allowed us to estimate the progression of the pathology and its correlates in cortical plasticity. The results indicate that visual evoked responses are robust in both Tau− and Tau+ mice in both age groups. However, Tau+ animals show reduced visual cortical plasticity at short (tens of seconds) and long (days) timescales. At 8 months of age, visual plasticity is reduced in Tau− animals, and practically abolished in Tau+ animals, potentially indicating a combined effect of age and tauopathy. In addition, we found an absence of behavioural responses to novel visual stimuli in Tau+ animals. Overall, these data indicate that tau-associated neurodegeneration has an impact on both long and short-term visual plasticity, and their potential behavioural correlates.

### Visual cortical plasticity in neurodegeneration

Previous work has found limited changes in visual cortical activity during neurodegeneration. Mice that overexpress amyloid-β precursor protein (APP) show robust orientation and direction selectivity in V1 (Grienberger et al., 2012; Liebscher et al., 2016). rTg4510 mice also show robust orientation tuning in V1, even at advanced stages of neurodegeneration (8-10 months) (Kuchibhotla et al., 2014). Similarly, we have shown that naive visual evoked responses in V1 of rTg4510 mice are largely unaffected in Tau+ animals. The basic functional properties of visual cortical neurons are, however, often explained by the functional properties of thalamocortical relay cells and the pattern of their cortical projections (Priebe, 2016). These basic functional properties may therefore be resilient, because the thalamus is largely unaffected in many models of neurodegeneration, including the rTg4510 model. In contrast, degeneration is more likely to have an earlier effect on functional properties that depend on cortical cellular mechanisms, and the balance of intra-cortical excitation and inhibition (Rubin et al., 2015). We therefore hypothesised that visual plasticity would be a more sensitive assay of the impact of degeneration on neuronal function.

We monitored visual plasticity at short- and long timescales. Stimulus response potentiation (SRP) is a form of long-term plasticity that is dependent on parvalbumin-positive interneuron activity (Kaplan et al., 2016), and is thought to co-opt similar pathways to thalamocortical LTP (Cooke and Bear, 2010), including synaptic plasticity, NMDA receptor activation and increased AMPA receptor trafficking (Cooke et al., 2015; Frenkel et al., 2006). Some of these circuits, particularly parvalbumin interneurons, are also thought to be important in ocular dominance plasticity (Hensch, 2005; Hooks and Chen, 2020), so our observation of reduced SRP in Tau+ animals may be consistent with disruption of ocular dominance plasticity in the visual cortex of APP and PS1 mouse models (Maya-Vetencourt et al., 2014; William et al., 2012). Our experimental design, however, allows for measurements at multiple timescales, and we found concomitant changes at short and long time scales of plasticity. Short-term visual plasticity, usually known as adaptation, is linked to transient changes in the responsivity of synapses (Regehr, 2012) or post-synaptic mechanisms related to spike frequency adaptation (Sanchez-Vives et al., 2000). Our results therefore suggest that even early stages of neurodegeneration have an impact on both synaptic mechanisms and intracellular trafficking in V1. We note that because we measured the local field potential, our measurements are likely to reflect the pooled signal of excitatory and inhibitory synapses (Einevoll et al., 2013; Kim et al., 2019). Spiking activity in individual neurons, which depends on idiosyncratic and finely balanced excitation and inhibition, may show variable effects.

We have shown that a simple visual paradigm allows easy characterisation of the impact of neurodegeneration on plasticity in awake animals, both at short (tens of seconds) and longer (days) time scales. We have also shown that plasticity over longer time scales is markedly reduced in older animals, consistent with age-related reductions in ocular dominance plasticity in mice that have been documented previously (Lehmann and Lowel, 2008; Lehmann et al., 2012). In humans, repetitive visual stimulation produces a lasting enhancement of VEPs as measured by EEG (Clapp et al., 2012; Teyler et al., 2005), similar to SRP. This potentiation has been shown to be impaired in disorders that are thought to be associated with hypofunction of the NMDA receptor, like schizophrenia (Cavus et al., 2012). AD patients also show a deficit in NMDAR-dependent forms of cortical plasticity (Battaglia et al., 2007). Assessing whether VEP potentiation is impaired in individuals with or at risk of AD, or in normal ageing, could be a subject for future studies. If true, it could provide a framework to assess disease progression in these patients that could be less invasive and cheaper than other biomarkers.

### Plasticity in rTg4510 mouse model of tauopathy

Previous work in rTg4510 mice has shown a link between neurodegeneration and plasticity in high-level cortical circuits. rTg4510 mice show impaired long-term depression in perirhinal cortex synapses, which may underlie defects in long-term recognition memory (Scullion et al., 2019). Impairments in LTP are also observed in the hippocampus (Booth et al., 2016; Gelman et al., 2018; Hoover et al., 2010) and frontal cortex (Crimins et al., 2012) of rTg4510 (and APP/PS1) mice, including changes in intrinsic membrane properties, depolarisation of the resting potential, increased excitability and changes in dynamics. These cellular changes are accompanied by disturbed oscillations in the LFP and disordering of ‘place cells’ in hippocampus (Cheng and Ji, 2013; Ciupek et al., 2015; Witton et al., 2016). These deficits in both short- and medium-term neuronal plasticity are not simply a result of accumulation of insoluble NFT, and arise before pronounced changes in cellular morphology (Crimins et al., 2012; Jackson et al., 2020; Rocher et al., 2010). Some of the functional effects may be explained by increased synaptic instability as a result of synaptic density reduction and increased dendritic spine turnover (Jackson et al., 2017) or may reflect mislocalization of soluble tau to dendrites (Zhao et al., 2016). The results we observed here are in agreement with these observations: Tau+ animals show reduced visual cortical plasticity early in neurodegeneration.

We note that retinal and optic nerve atrophy has been reported in late-stages (ca. 7.5 months) of rTg4510 neurodegeneration (Harrison et al., 2019). These changes in retinal structure may be expected to reduce the visual cortical response as measured by the VEP. However, we found that VEP amplitude was robust, even in 8-months old Tau+ animals. Tau+ mice also have altered circadian rhythms, with increased periods of wakefulness, less time in nonrapid-eye-movement sleep (Holton et al., 2020), and altered sharp-wave ripple dynamics (Witton et al., 2016). These changes may contribute to the changes in long-term visual plasticity that we observed as sleep is important for SRP in mouse V1 (Aton et al., 2014; Cooke and Bear, 2010, 2012; Ji and Wilson, 2007).

Recent work (Gamache et al., 2019) shows that in the rTg4510 mouse model, a fragment of the *Fgf14* gene was replaced by the insertion of the P301L transgene (tau-Tg). This genetic disruption may influence the progress of neuronal loss and behavioural abnormalities alone or in combination with the expression of the mutant tau. Our experiments mitigate these potential offsite effects by comparing Tau− and Tau+ mice, that differ in mutant tau expression but not genotype. In addition, our comparisons of Tau− with WT littermates reveal similar long-term plasticity and visual evoked behaviour. Interestingly, older (8 m) Tau− mice showed reduced adaptation, or short-term plasticity but WT animals did not. Adaptation effects are generally thought to be preserved with age in humans (Elliott et al., 2007; Elliott et al., 2012). It is possible that reduced adaptation in Tau− animals could be an effect of the *Fgf14* deletion or it could be an effect of tau accumulation before the onset of the doxycycline treatment (which started at 2 months). If the latter is true, this would render adaptation as a more sensitive assay to assess functional changes in neurodegeneration in older animals.

Previous studies have reported alterations in rTg4510 mouse behaviour: Tau+, and to a lesser extent, Tau− mice tend to show hyperactivity and motor stereotypy, whilst maintaining good motor coordination (Blackmore et al., 2017; Cook et al., 2014; Holton et al., 2020; Jul et al., 2016; Wes et al., 2014). These mice do not appear to respond to novelty, and have impaired spatial working memory (Blackmore et al., 2017; Wes et al., 2014). In our experiments on head-fixed animals, Tau+ animals reacted less to the appearance of a new visual stimulus than did their Tau− or WT counterparts, consistent with reduced novelty responses. This insensitivity to novelty may partly reflect a lack of arousal or a deficit in attention, which may in turn indicate deficiencies in noradrenaline circuits that appear critical for these processes (Sara, 2009). The CamKIIa promoter that drives transgene expression in the rTg4510 mouse is likely to be expressed in locus coeruleus (Glennon et al., 2019), and it is therefore possible that the noradrenalin input to cortex is disrupted in Tau+ mice.

### Conclusions

Plasticity is a hallmark of neuronal function, important for learning and memory. Neurodegenerative diseases like AD not only disrupt neuronal structure and function, but erode the flexibility of neurons and circuits. Failure of synaptic plasticity is assumed to occur early in the course of AD. We verified the effect of tauopathy on a form of sensory learning and memory in V1 of mouse. We found impaired visual plasticity even at early stages of neurodegeneration, before substantial neuronal loss occurs. Our measurements offer a simple and direct measurement of plasticity in degenerating brain circuits, and a potential target for understanding, detecting and tracking that neurodegeneration.

## Acknowledgements

We thank Francesca Cacucci (F. C.) for comments on the manuscript. We thank Zeshan Ahmed, and Anthony Blockeel for useful advice and discussions. This work was supported by the Medical Research Council grant (R023808) to S.G.S., A.B.S. and F.C., a Biotechnology and Biological Sciences Research Council grant (R004765) to S.G.S. and A.B.S., a Sir Henry Dale Fellowship from the Wellcome Trust and Royal Society (200501) to A.B.S, and an International Collaboration Award to S.G.S (with Adam Kohn) from the Stavros Niarchos Foundation / Research to Prevent Blindness.

## Author Contributions

Conceptualization, A.P., A.B.S., and S.G.S.; Methodology, A.P., F.R.R., A.B.S., and S.G.S.; Investigation, A.P., F.R.R., and J.H.; Validation, Formal Analysis, Data Curation, A.P., F.R.R., S.G.S.; Writing – Original Draft, A.P. and F.R.R.; Writing – Review & Editing, A.P., F.R.R., A.B.S., and S.G.S.; Visualization, A.P., F.R.R., A.B.S., and S.G.S.; Funding Acquisition, A.B.S., and S.G.S.; Resources, A.B.S., K.P. and S.G.S.; Supervision, A.B.S., and S.G.S.

## Methods

### Animal Experiments

All experiments were performed in accordance with the Animals (Scientific Procedures) Act 1986 (United Kingdom) and Home Office (United Kingdom) approved project and personal licenses. The experiments were approved by the University College London Animal Welfare Ethical Review Board under Project License 70/8637.

The generation of rTg4510 transgenic mice was performed as described previously (Blackmore et al., 2017; Santacruz et al., 2005). A total of 50 male transgenic mice and 16 wild-type (WT) littermates were obtained at approximately 7 weeks of age from Eli Lilly and Company (Windlesham, UK) via Envigo (Loughborough, UK). At 8 weeks of age, to suppress Tau expression, 25 of the 50 transgenic mice were treated with Doxycycline, which included four 10mg/kg bolus oral doses of doxycycline (Sigma) in 5% glucose solution by oral gavage across 4 days, followed by *ad libitum* access to Teklad base diet containing 200ppm doxycycline (Envigo) for the duration of the experiment. The mice in this group were designated ‘Tau−’ animals. The remaining 25 animals, designated as ‘Tau+’, and the WT animals received 4 oral doses of the vehicle (5% glucose) and had *ad libitum* access to standard feed for the duration of the experiment. All animals also had *ad libitum* access to water. Mice were subsequently divided into two cohorts. One cohort was tested at approximately 5 months (22-26 weeks, 12 Tau−, 12 Tau+, 6 WT), and another cohort was tested at approximately 8 months of age (31-35 weeks, 13 Tau−, 13 Tau+, 10 WT). Mice were group housed to a maximum of 5 individuals per cage until 3 days before surgery, when they were separated into individual cages. All animals were kept under a 12-hour light/dark cycle, and both behavioural and electrophysiological recordings were carried out during the dark phase of the cycle.

### Surgery

Mice were anaesthetised for surgery with 3% isoflurane in O_2_. Preoperative analgesia (Carprieve, 5mg/kg) was given subcutaneously and lubricant ophthalmic ointment was applied. Anaesthesia was maintained with 1-2% isoflurane in O_2_ and the depth was monitored by absence of pinch-withdrawal reflex and breathing rate. The body temperature was maintained using a heating blanket. A small craniotomy hole (<1mm^2^) was made over the right primary visual cortex (2.8mm lateral and 0.5mm anterior from lambda) and a chronic local field potential (LFP) recording electrode (Bear lab chronic microelectrode Monopolar 30070, FHC, USA) was implanted approximately 400-450μm below the cortical surface. A ground screw was implanted over the left prefrontal cortex and a custom-built stainless-steel metal plate was affixed on the skull. Dental cement (Super-Bond C&B, Sun Medical) was used to cover the skull, ground screw and metal plate, enclosing and stabilizing the electrode. Analgesic treatment (Metacam, Boehringer Ingelheim, 1mg/kg) mixed in condensed milk was provided orally for three days after the surgery. Mice recovered for at least 7 days before the first recording session.

### Visual Stimulus Presentation & Experimental Setup

Visual stimuli consisted of full-field, 100% contrast sinusoidal gratings generated using BonVision (Lopes et al., 2020), presented on a γ-corrected computer monitor (Iiyama ProLite EE1890SD). The gratings were presented in a circular aperture with hard edges, outside of which the monitor was held at the mean luminance (‘grey screen’). The grating was oriented at either −45° or 45° from vertical, and reversed contrast (flickered) at a frequency of 1.95Hz. The display was placed 15cm from – and normal to – the mouse and centred on the left visual field. The stimulus was warped to maintain visual angle across the monitor.

One week after the surgery, mice were placed on a styrofoam wheel with a grey screen and habituated over 5 days to the experimental set-up by progressively increasing the time spent head-fixed, from ~5 mins to 30 mins. Mice were allowed to run on the wheel, and their speed was recorded using a rotary encoder. Pupils were imaged using an infrared camera camera (DMK 22BUC03, ImagingSource; 30 Hz) focused on the left eye through a zoom lens (Computar MLH-10X Macro Zoom Lens), and acquired by the same computer that presented the visual stimulus. Pupil estimates (position, diameter) were tracked online using custom routines in Bonsai. At the beginning of each recording session, mice were presented with a grey screen for 3-5min. On the first day of recordings, mice were presented with 5 blocks of a grating oriented at 45° and 5 blocks of a grating oriented at −45°, alternating between the two. Each block consisted of 200 continuous reversals, and were separated by 30s, during which the monitor was held at the mean luminance. For 6 animals (2 WT, 2 Tau− and 2 Tau+), each block consisted of 400 phase reversals. On days 2-8, mice were presented with 10 blocks of a single oriented grating (familiar stimulus), that was either 45° or −45°, randomly assigned for each animal (counterbalanced between groups). The last day of recordings, day 9, was similar to day 1. For days 1 and 9, whether the first block presented would be the familiar or the unfamiliar stimulus (presented only on the first and last day of recordings), was randomly assigned for each animal.

### Neural Recordings

Signals from the recording electrode were acquired, digitised and filtered using an OpenEphys acquisition board connected to a different computer from that used to generate the visual stimulus. The electrophysiological and rotary wheel signals were sampled at 30kHz. These data were synchronised with pupil video recordings and visual stimulus via the signal of a photodiode (PDA25K2, Thorlabs, Inc., USA) that monitored timing pulses on a small corner of the monitor shielded from the animal.

### Data analysis

All data were analysed using custom software written in MATLAB (MathWorks). Neural and wheel data were filtered with an 8th order Chebyshev Type I lowpass filter and downsampled to 1kHz.

#### VEP analysis

VEPs were averaged across all phase reversals and blocks for each stimulus on each day. To estimate the VEP amplitude, the LFP signal was filtered using a second order bandpass filter with a 0.3Hz low cut and 50Hz high cut frequency. The negative trough was defined as the minimum value within the first 150ms after stimulus reversal, and the positive peak was defined as the maximum value within the first 250ms. The amplitude was defined as the difference between this trough and peak.

#### Pupil data

Eye blinks were removed by identifying any points that were two times above the variance of the eye position. Removed values were replaced using nearest neighbour interpolation. Pupil area (in pixels) was converted to mm^2^ by multiplying with the square of the camera resolution (in mm/pixel). Responses were then normalised to the average pupil area in the two minutes before the stimulus onset, where animals were viewing a grey screen.

#### Wheel data

We estimated the speed and direction of the rotating wheel using a quadrature encoder. Rotations were converted to speed by multiplying with the wheel circumference and dividing by the encoder resolution. Speed was then smoothed using a gaussian filter with a 50ms window.

#### Regression

We fitted an elastic net regularization model to predict the VEP amplitude for each animal using days, block number within each day, movement speed and pupil area as regressors. Regressors and predicted value were normalized to range between 0 and 1 before fitting. We explored the entire regularisation path by using N values (<=10,000) of the regularisation hyperparameter, fixing the ratio of the λ1 and λ2-norm regularisers to 2. The impact M of each regressor r in predicting the VEP amplitude was calculated as:

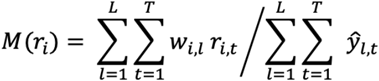

Where *l* is the regularization hyperparameter, *t* denotes each time there is a stimulus reversal (*t* = 1: *number of reversal each day* * *number of days*), *w* is the learned weight, *r_i_* denotes each respective regressor (where *i* is either days, block number, speed, pupil area) and *y* is the predicted value of the elastic net model. Note that if two or more regressors are correlated, the regularization model can assign weights to one or both regressors.

#### Visual adaptation

To estimate the adaptation effect for each animal, for each day, the VEP amplitude was calculated within a block using a step of 20 reversals. The mean amplitude was subtracted from each block and amplitudes were averaged over blocks 2-10. A decaying exponential function was fitted to the averaged data using least squares. The exponential time constant for each animal was fixed to τ=7.6 reversals. This value was calculated by fitting the exponential to the mean trace obtained by averaging over all days and all animals.

### Brain samples

Mice were euthanised by overdose injection of pentobarbital (intraperitoneal) and perfused with phosphate-buffered saline (PBS). Brains were removed, weighed, and the right hemisphere was fixed in 10% buffered formalin until processed (7-13 months) for immunohistochemistry pathology assessment.

### Histopathology

Immunohistochemistry was performed for all mice. The brains were immersed in PBS that contained 30% sucrose, and subsequently cut into 40μm parasagittal sections on a cryostat (Leica CM1520). Antigen retrieval was achieved through heating sections in citrate buffer (pH 6.0; Vector Labs) in an oven at 60°C overnight. Slides were treated with 0.3% hydrogen peroxide solution (3% in distilled water) for 10min to eliminate endogenous peroxidase activity, and subsequently washed with PBS with 0.5% Triton X-100. Immunohistochemistry was performed using a primary antibody for tau phosphorylated at serine 202 (mouse monoclonal AT8, 1:1000, Thermo Fisher Scientific). The Mouse on Mouse (MOM) Detection Kit (Vector Labs, BMK-2202) was used for staining, with buffers prepared as described in standard protocol supplied with the kit. After rinsing, slides were treated for 5 min with the chromogen 3,3’-diaminobenzidine (DAB; Vector Laboratories, SK-4105) to allow visualisation. The slides were then coverslipped with Shandon ClearVue Mountant XYL (Thermofisher) and digitised using a Leica Microscope (DMi8 S) coupled with a Leica camera (DFC7000 GT; Leica Microsystems). The Fiji image processing package was used to view the digitized tissue sections (Schindelin et al., 2012). At least 3 regions of interest (ROIs) of the same size were selected from approximately the same brain location in V1 and hippocampus for each animal. In cases where V1 sections were fragmented (n=12), additional ROIs were selected from elsewhere in the cortex. To assess the tau burden, a normal distribution was fit to the image of each ROI and the mean and standard deviation were obtained and averaged over all ROIs for each animal (Fig. S1). To compare the tau burden in V1 and hippocampus, the mean values obtained from the fitted distributions were zscored and sign inverted (Fig. 1). These analyses were performed in a blinded fashion.

### Statistical analyses

All data are presented as a mean ± standard error of the mean (SEM). Statistical comparisons were performed in MATLAB and SPSS (IBM). A two-way or a repeated-measures ANOVA was applied for all comparisons. p < 0.05 was used as the significance threshold. Exact p values are given.

## Supplementary Figures

**Figure S1:**
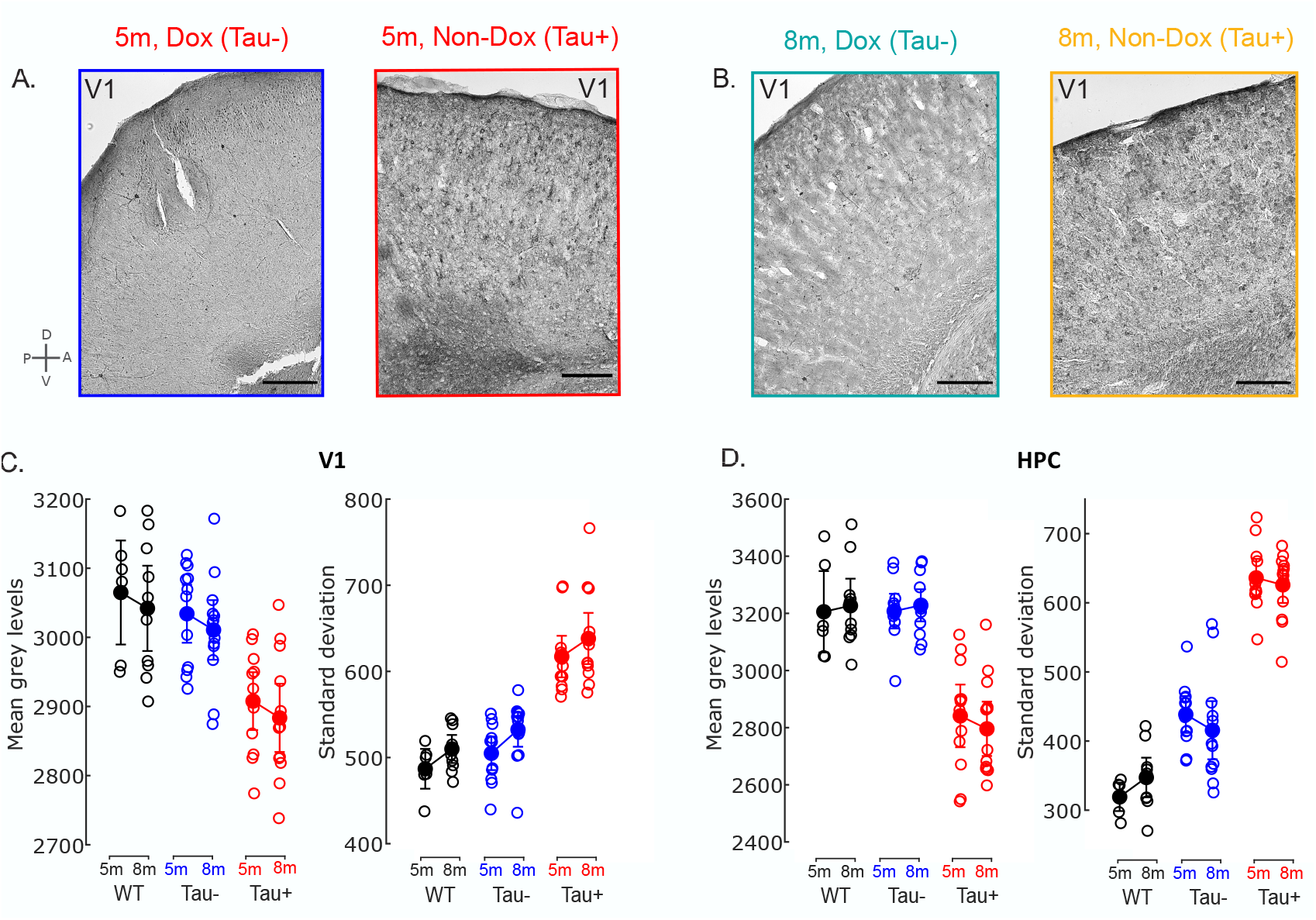
Immunohistochemical profiling of 5- and 8-month old rTg4510 mice. **A-B.** Representative immunohistochemical images from the V1 of Tau− mice receiving doxycycline (Dox) treatment and Tau+ mice (Non-Dox) at 5 months old (**A**) and 8 months old (**B**). **C.** Average of the mean grey levels (left) and standard deviation (right) of ROIs selected in V1 for each animal (Methods). Data are represented as mean±2*SEM. **D.** Same as in C. for hippocampal ROIs. Tau+ animals were characterised by a smaller mean and larger standard deviation compared to Tau− and WT animals, confirming that the doxycycline treatment was successful.

**Figure S2:**
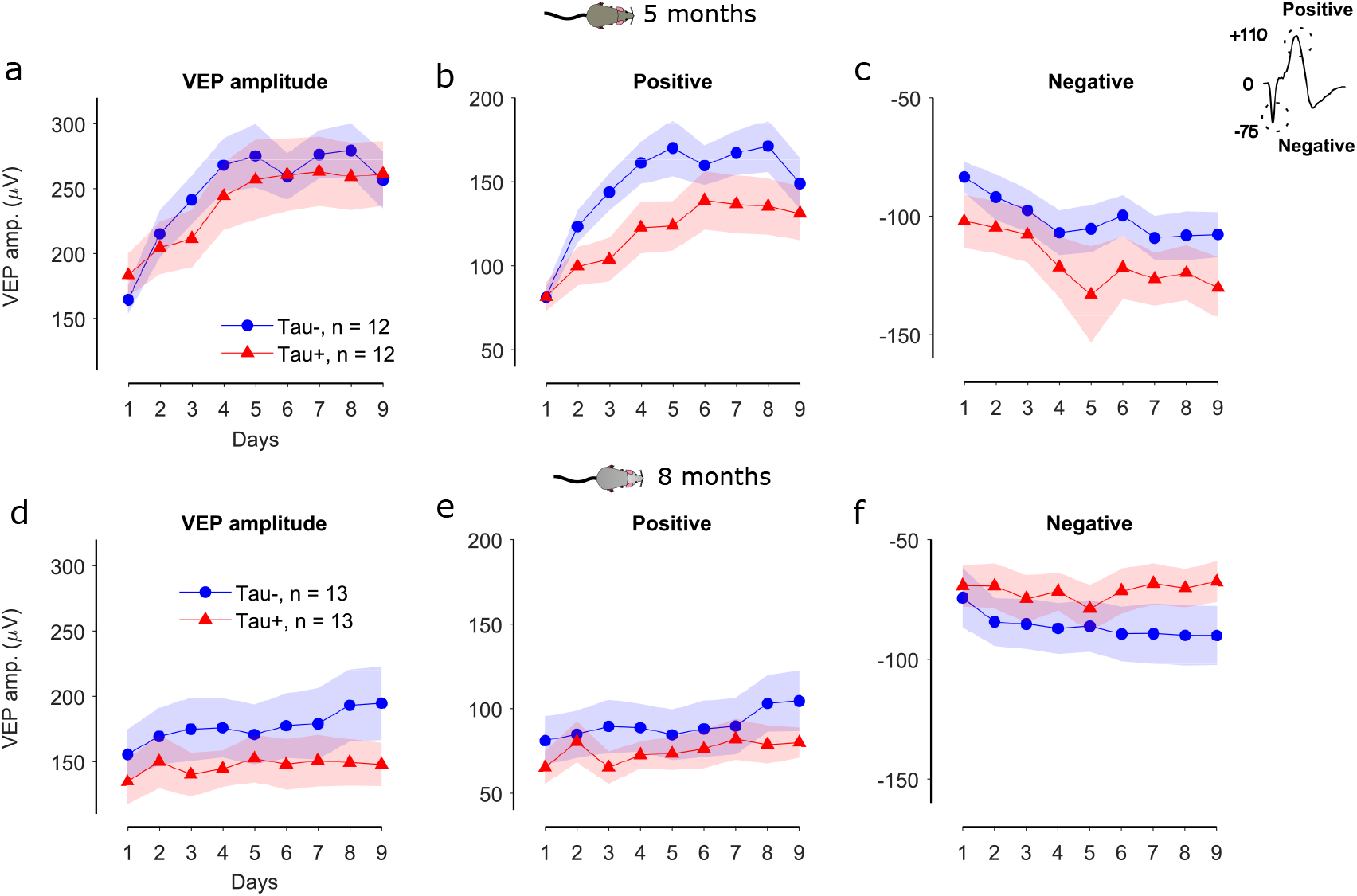
Differences in visual plasticity between Tau+ and Tau− animals are attributed mainly to the positive deflection of the VEP signal. **a.** Average VEP amplitude, defined as the difference between the positive and negative peaks of the VEP signal, across days. **b.** Difference between the positive deflection of the VEP signal and the baseline as a function of days, averaged across Tau+ and Tau− animals. **c.** Difference between the negative deflection of the VEP signal and the baseline as a function of days, averaged across the groups of animals. **d-f.** Same for 8-month old animals.

**Figure S3:**
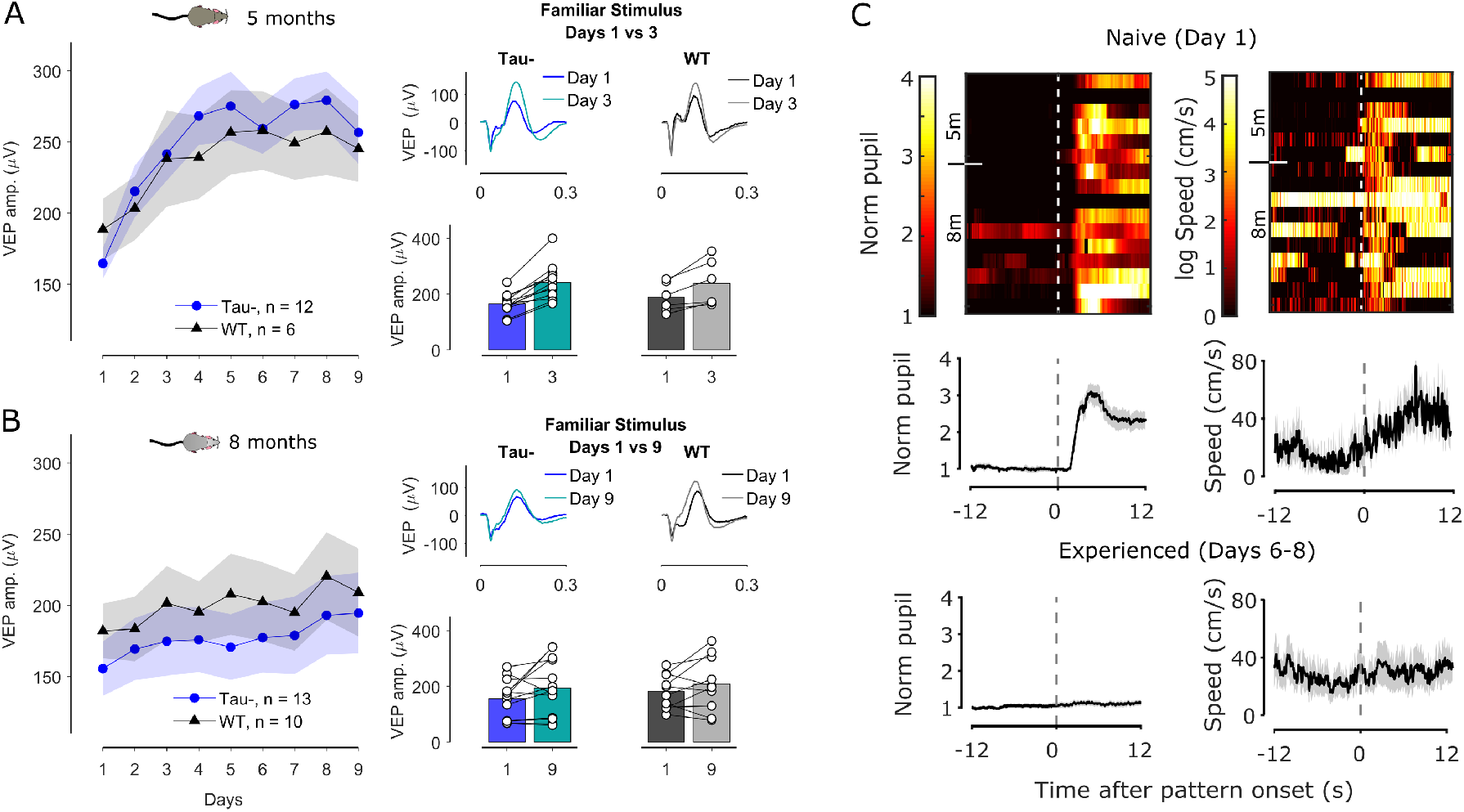
VEP and behavioural responses are similar in Tau− and Wild Type (WT) mice. **A.** Average VEP amplitude as a function of days for 5-month old Tau− mice (blue) and WT littermates (black). The panels on the right show the average VEP signal on top and the VEP amplitude of individual animals on the bottom for days 1 and 3. VEPs significantly increased by day 3 relative to day 1 for the familiar stimulus for both Tau− and WT animals. **B.** Average VEP amplitude as a function of days for 8-month old Tau− mice (blue) and WT littermates (black). The panels on the right show the average VEP signal and the VEP amplitude of individual animals for days 1 and 9. WT mice had on average a larger VEP amplitude than Tau− mice but the VEPs increased at a similar rate as a function of days. The rate and magnitude of potentiation was reduced compared to 5-month old animals. **C.** Left: Images and average responses of the normalized pupil to the onset of the stimulus of the first block presented for naive (top) and experienced (bottom) WT mice. Right: Images and average responses of the movement speed.

**Figure S4:**
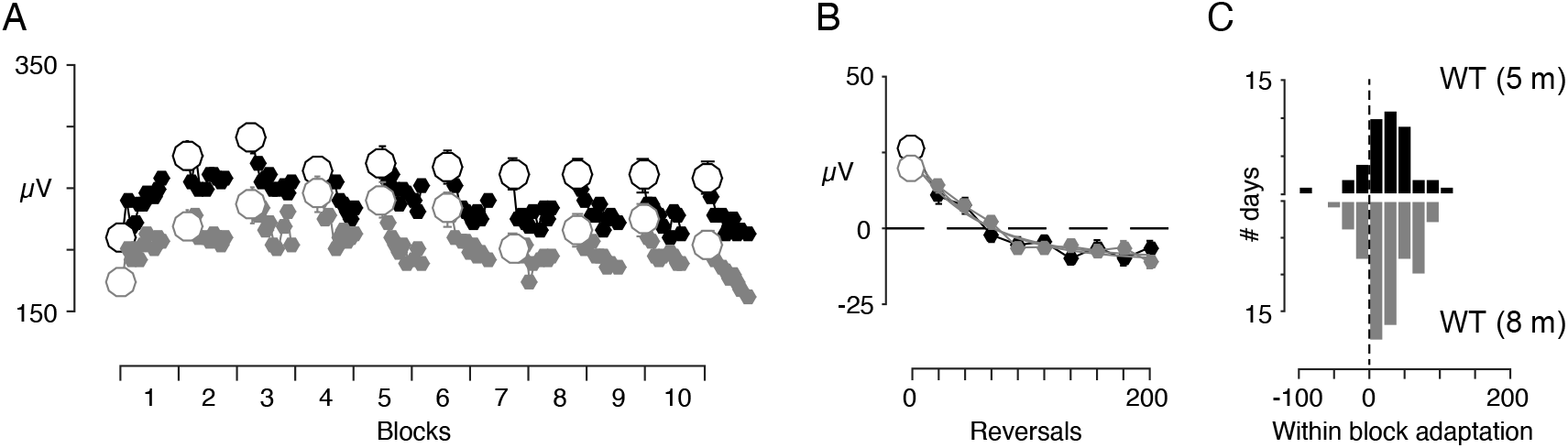
Adaptation in WT animals does not reduce with age (related to Figure 4). **A.** Average VEP amplitude as a function of block number for 5-month old WT mice (black) and 8-month old WT mice (grey). The VEP amplitude was calculated from non-overlapping averages of 20 reversals. With the notable exception of the first block, VEPs showed a reduction of responses within each block, consistent with classic sensory adaptation effects. **B.** Average VEP responses for blocks 2-10 on days 2-8. **C.** Histograms of the fitted amplitudes showing no significant differences with age.

**Figure S5:**
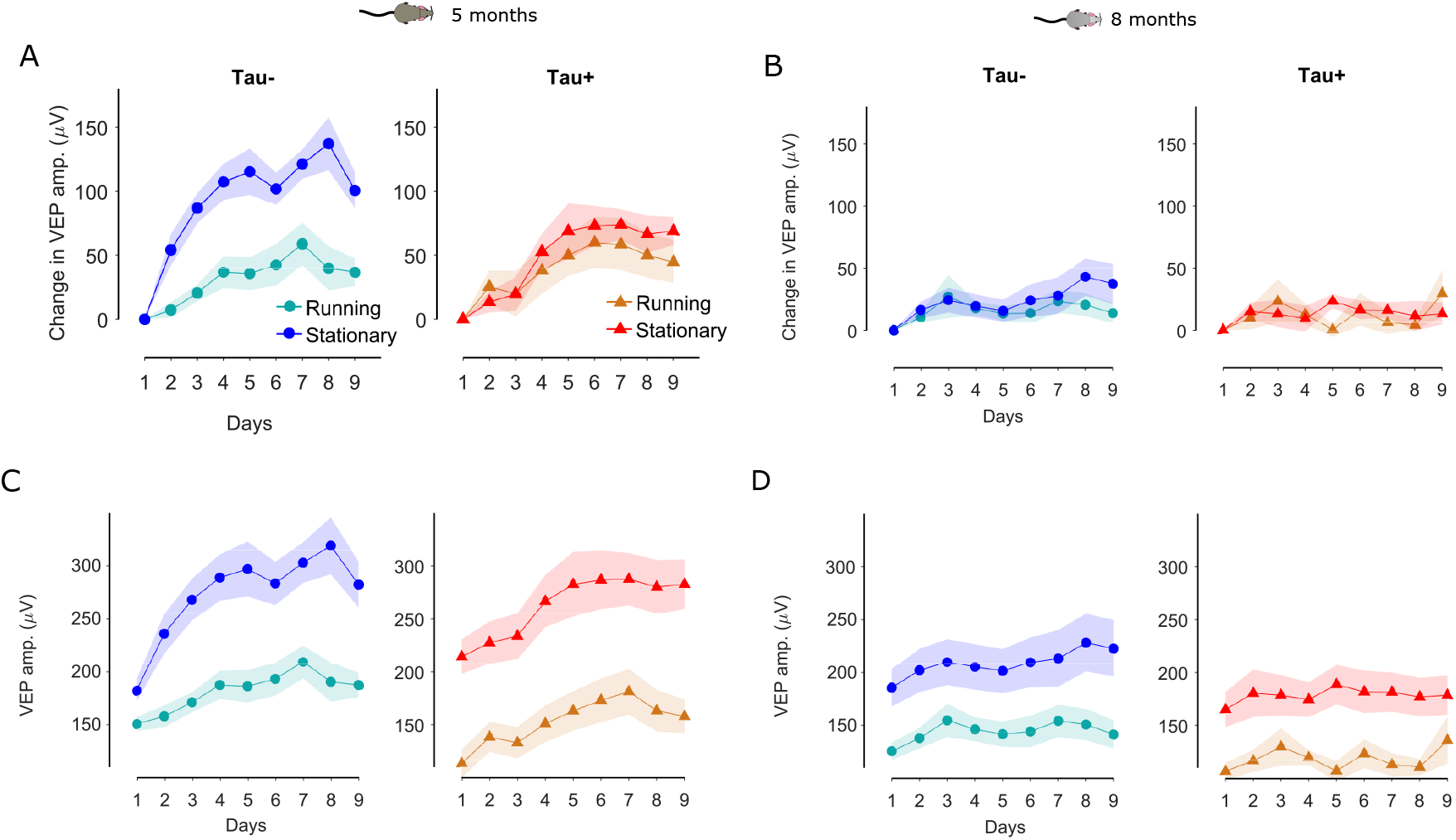
Running reduces the VEP amplitude but does not affect stimulus-response potentiation in either Tau+ or Tau− animals. **A.** Difference in the VEP amplitude from day 1 for the familiar stimulus for Tau− (left) and Tau+ (right) 5-month old animals over the course of days calculated considering only stationary or running epochs. **B.** Same as in A for 8-month old animals. **C.** Unnormalized VEP amplitude as a function of days for 5-month old Tau− (left) and Tau+ (right) mice calculated considering only stationary or running epochs. **D.** Same as in C for 8-month old animals.

**Figure S6:**
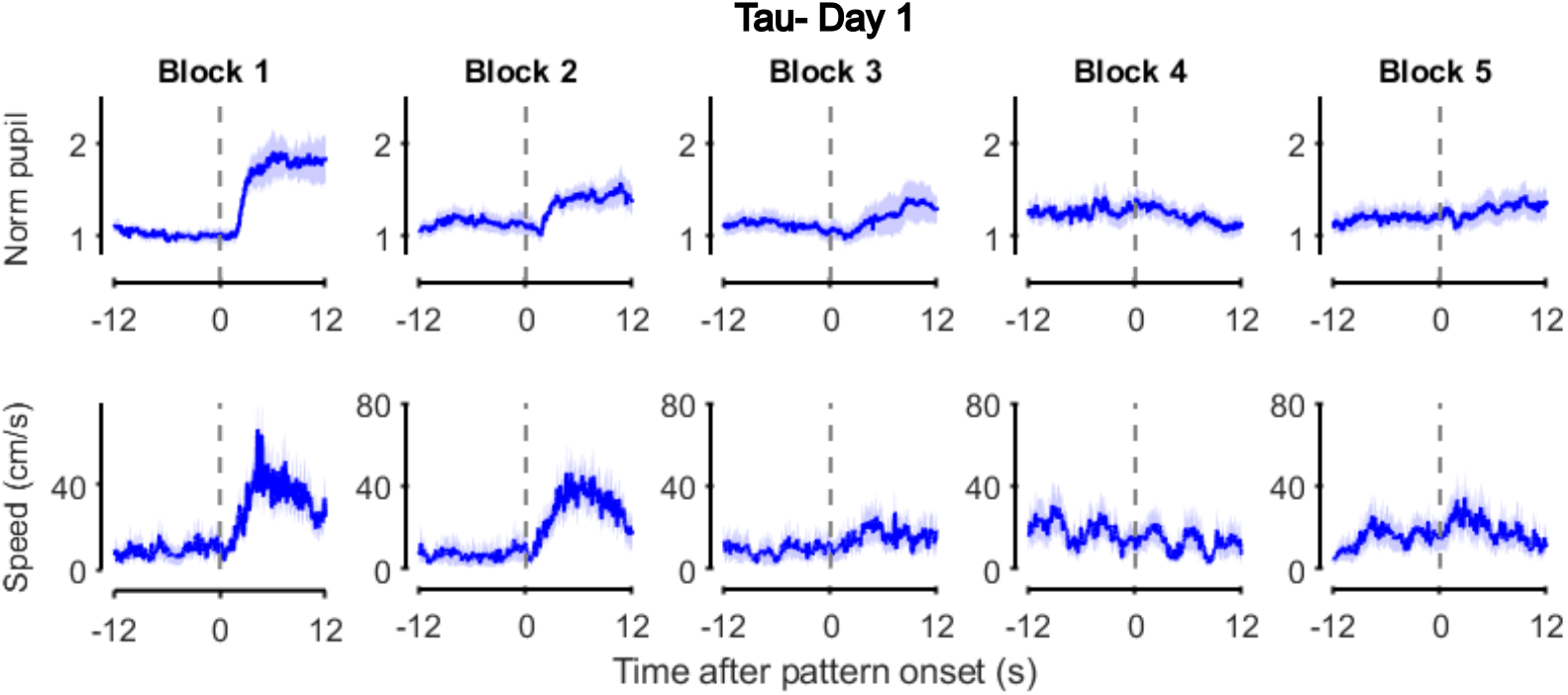
Visual evoked behaviour habituates quickly in Tau− animals. Mean (±SEM) normalized pupil responses (top) and movement speed (bottom) to the onset of the stimulus during the first five blocks of presentation for naive Tau− animals (day 1).

## References

Aton, S.J., Suresh, A., Broussard, C., and Frank, M.G. (2014). Sleep promotes cortical response potentiation following visual experience. Sleep 37, 1163–1170.

Battaglia, F., Wang, H.Y., Ghilardi, M.F., Gashi, E., Quartarone, A., Friedman, E., and Nixon, R.A. (2007). Cortical plasticity in Alzheimer’s disease in humans and rodents. Biol Psychiatry 62, 1405–1412.

Blackmore, T., Meftah, S., Murray, T.K., Craig, P.J., Blockeel, A., Phillips, K., Eastwood, B., O’Neill, M.J., Marston, H., Ahmed, Z., et al. (2017). Tracking progressive pathological and functional decline in the rTg4510 mouse model of tauopathy. Alzheimers Res Ther 9, 77.

Booth, C.A., Witton, J., Nowacki, J., Tsaneva-Atanasova, K., Jones, M.W., Randall, A.D., and Brown, J.T. (2016). Altered Intrinsic Pyramidal Neuron Properties and Pathway-Specific Synaptic Dysfunction Underlie Aberrant Hippocampal Network Function in a Mouse Model of Tauopathy. J Neurosci 36, 350–363.

Cavus, I., Reinhart, R.M., Roach, B.J., Gueorguieva, R., Teyler, T.J., Clapp, W.C., Ford, J.M., Krystal, J.H., and Mathalon, D.H. (2012). Impaired visual cortical plasticity in schizophrenia. Biol Psychiatry 71, 512–520.

Cheng, J., and Ji, D. (2013). Rigid firing sequences undermine spatial memory codes in a neurodegenerative mouse model. Elife 2, e00647.

Ciupek, S.M., Cheng, J., Ali, Y.O., Lu, H.C., and Ji, D. (2015). Progressive functional impairments of hippocampal neurons in a tauopathy mouse model. J Neurosci 35, 8118–8131.

Clapp, W.C., Hamm, J.P., Kirk, I.J., and Teyler, T.J. (2012). Translating long-term potentiation from animals to humans: a novel method for noninvasive assessment of cortical plasticity. Biol Psychiatry 71, 496–502.

Cook, C., Dunmore, J.H., Murray, M.E., Scheffel, K., Shukoor, N., Tong, J., Castanedes-Casey, M., Phillips, V., Rousseau, L., Penuliar, M.S., et al. (2014). Severe amygdala dysfunction in a MAPT transgenic mouse model of frontotemporal dementia. Neurobiol Aging 35, 1769–1777.

Cooke, S.F., and Bear, M.F. (2010). Visual experience induces long-term potentiation in the primary visual cortex. J Neurosci 30, 16304–16313.

Cooke, S.F., and Bear, M.F. (2012). Stimulus-selective response plasticity in the visual cortex: an assay for the assessment of pathophysiology and treatment of cognitive impairment associated with psychiatric disorders. Biol Psychiatry 71, 487–495.

Cooke, S.F., Komorowski, R.W., Kaplan, E.S., Gavornik, J.P., and Bear, M.F. (2015). Visual recognition memory, manifested as long-term habituation, requires synaptic plasticity in V1. Nat Neurosci 18, 262–271.

Crimins, J.L., Rocher, A.B., and Luebke, J.I. (2012). Electrophysiological changes precede morphological changes to frontal cortical pyramidal neurons in the rTg4510 mouse model of progressive tauopathy. Acta Neuropathol 124, 777–795.

Crutch, S.J., Yong, K.X., and Shakespeare, T.J. (2016). Looking but not seeing: Recent perspectives on posterior cortical atrophy. Current Directions in Psychological Science 25, 251–260.

Einevoll, G.T., Kayser, C., Logothetis, N.K., and Panzeri, S. (2013). Modelling and analysis of local field potentials for studying the function of cortical circuits. Nat Rev Neurosci 14, 770–785.

Elliott, S.L., Hardy, J.L., Webster, M.A., and Werner, J.S. (2007). Aging and blur adaptation. J Vis 7, 8.

Elliott, S.L., Werner, J.S., and Webster, M.A. (2012). Individual and age-related variation in chromatic contrast adaptation. J Vis 12, 11.

Frenkel, M.Y., Sawtell, N.B., Diogo, A.C., Yoon, B., Neve, R.L., and Bear, M.F. (2006). Instructive effect of visual experience in mouse visual cortex. Neuron 51, 339–349.

Gamache, J., Benzow, K., Forster, C., Kemper, L., Hlynialuk, C., Furrow, E., Ashe, K.H., and Koob, M.D. (2019). Factors other than hTau overexpression that contribute to tauopathy-like phenotype in rTg4510 mice. Nat Commun 10, 2479.

Gelman, S., Palma, J., Tombaugh, G., and Ghavami, A. (2018). Differences in Synaptic Dysfunction Between rTg4510 and APP/PS1 Mouse Models of Alzheimer’s Disease. J Alzheimers Dis 61, 195–208.

Glennon, E., Carcea, I., Martins, A.R.O., Multani, J., Shehu, I., Svirsky, M.A., and Froemke, R.C. (2019). Locus coeruleus activation accelerates perceptual learning. Brain Res 1709, 39–49.

Grienberger, C., Rochefort, N.L., Adelsberger, H., Henning, H.A., Hill, D.N., Reichwald, J., Staufenbiel, M., and Konnerth, A. (2012). Staged decline of neuronal function in vivo in an animal model of Alzheimer’s disease. Nat Commun 3, 774.

Grundke-Iqbal, I., Iqbal, K., Tung, Y.C., Quinlan, M., Wisniewski, H.M., and Binder, L.I. (1986). Abnormal phosphorylation of the microtubule-associated protein tau (tau) in Alzheimer cytoskeletal pathology. Proc Natl Acad Sci U S A 83, 4913–4917.

Harrison, I.F., Whitaker, R., Bertelli, P.M., O’Callaghan, J.M., Csincsik, L., Bocchetta, M., Ma, D., Fisher, A., Ahmed, Z., Murray, T.K., et al. (2019). Optic nerve thinning and neurosensory retinal degeneration in the rTg4510 mouse model of frontotemporal dementia. Acta Neuropathol Commun 7, 4.

Helboe, L., Egebjerg, J., Barkholt, P., and Volbracht, C. (2017). Early depletion of CA1 neurons and late neurodegeneration in a mouse tauopathy model. Brain Res 1665, 22–35.

Hensch, T.K. (2005). Critical period plasticity in local cortical circuits. Nat Rev Neurosci 6, 877–888.

Hof, P.R., Bouras, C., Constantinidis, J., and Morrison, J.H. (1990). Selective disconnection of specific visual association pathways in cases of Alzheimer’s disease presenting with Balint’s syndrome. J Neuropathol Exp Neurol 49, 168–184.

Holton, C.M., Hanley, N., Shanks, E., Oxley, P., McCarthy, A., Eastwood, B.J., Murray, T.K., Nickerson, A., and Wafford, K.A. (2020). Longitudinal changes in EEG power, sleep cycles and behaviour in a tau model of neurodegeneration. Alzheimers Res Ther 12, 84.

Hooks, B.M., and Chen, C. (2020). Circuitry Underlying Experience-Dependent Plasticity in the Mouse Visual System. Neuron 107, 986–987.

Hoover, B.R., Reed, M.N., Su, J., Penrod, R.D., Kotilinek, L.A., Grant, M.K., Pitstick, R., Carlson, G.A., Lanier, L.M., Yuan, L.L., et al. (2010). Tau mislocalization to dendritic spines mediates synaptic dysfunction independently of neurodegeneration. Neuron 68, 1067–1081.

Jackson, J.S., Johnson, J.D., Meftah, S., Murray, T.K., Ahmed, Z., Fasiolo, M., Hutton, M.L., Isaac, J.T.R., O’Neill, M.J., and Ashby, M.C. (2020). Differential aberrant structural synaptic plasticity in axons and dendrites ahead of their degeneration in tauopathy. bioRxiv, 2020.2004.2029.067629.

Jackson, J.S., Witton, J., Johnson, J.D., Ahmed, Z., Ward, M., Randall, A.D., Hutton, M.L., Isaac, J.T., O’Neill, M.J., and Ashby, M.C. (2017). Altered Synapse Stability in the Early Stages of Tauopathy. Cell Rep 18, 3063–3068.

Ji, D., and Wilson, M.A. (2007). Coordinated memory replay in the visual cortex and hippocampus during sleep. Nat Neurosci 10, 100–107.

Jul, P., Volbracht, C., de Jong, I.E., Helboe, L., Elvang, A.B., and Pedersen, J.T. (2016). Hyperactivity with Agitative-Like Behavior in a Mouse Tauopathy Model. J Alzheimers Dis 49, 783–795.

Kaplan, E.S., Cooke, S.F., Komorowski, R.W., Chubykin, A.A., Thomazeau, A., Khibnik, L.A., Gavornik, J.P., and Bear, M.F. (2016). Contrasting roles for parvalbumin-expressing inhibitory neurons in two forms of adult visual cortical plasticity. Elife 5.

Kim, T., Chaloner, F.A., Cooke, S.F., Harnett, M.T., and Bear, M.F. (2019). Opposing Somatic and Dendritic Expression of Stimulus-Selective Response Plasticity in Mouse Primary Visual Cortex. Front Cell Neurosci 13, 555.

Kuchibhotla, K.V., Wegmann, S., Kopeikina, K.J., Hawkes, J., Rudinskiy, N., Andermann, M.L., Spires-Jones, T.L., Bacskai, B.J., and Hyman, B.T. (2014). Neurofibrillary tangle-bearing neurons are functionally integrated in cortical circuits in vivo. Proc Natl Acad Sci U S A 111, 510–514.

Lehmann, K., and Lowel, S. (2008). Age-dependent ocular dominance plasticity in adult mice. PLoS One 3, e3120.

Lehmann, K., Schmidt, K.F., and Lowel, S. (2012). Vision and visual plasticity in ageing mice. Restor Neurol Neurosci 30, 161–178.

Liebscher, S., Keller, G.B., Goltstein, P.M., Bonhoeffer, T., and Hubener, M. (2016). Selective Persistence of Sensorimotor Mismatch Signals in Visual Cortex of Behaving Alzheimer’s Disease Mice. Curr Biol 26, 956–964.

Lopes, G., Farrell, K., Horrocks, E.A., Lee, C.-Y., Morimoto, M.M., Muzzu, T., Papanikolaou, A., Rodrigues, F.R., Wheatcroft, T., and Zucca, S. (2020). BonVision-an open-source software to create and control visual environments. BioRxiv.

Maya-Vetencourt, J.F., Carucci, N.M., Capsoni, S., and Cattaneo, A. (2014). Amyloid plaque-independent deficit of early postnatal visual cortical plasticity in the 5XFAD transgenic model of Alzheimer’s disease. J Alzheimers Dis 42, 103–107.

Mondragon-Rodriguez, S., Basurto-Islas, G., SantaMaria, I., Mena, R., Binder, L.I., Avila, J., Smith, M.A., Perry, G., and Garcia-Sierra, F. (2008). Cleavage and conformational changes of tau protein follow phosphorylation during Alzheimer’s disease. Int J Exp Pathol 89, 81–90.

Niell, C.M., and Stryker, M.P. (2010). Modulation of visual responses by behavioral state in mouse visual cortex. Neuron 65, 472–479.

Ossenkoppele, R., Schonhaut, D.R., Baker, S.L., O’Neil, J.P., Janabi, M., Ghosh, P.M., Santos, M., Miller, Z.A., Bettcher, B.M., Gorno-Tempini, M.L., et al. (2015). Tau, amyloid, and hypometabolism in a patient with posterior cortical atrophy. Ann Neurol 77, 338–342.

Priebe, N.J. (2016). Mechanisms of Orientation Selectivity in the Primary Visual Cortex. Annu Rev Vis Sci 2, 85–107.

Ramsden, M., Kotilinek, L., Forster, C., Paulson, J., McGowan, E., SantaCruz, K., Guimaraes, A., Yue, M., Lewis, J., Carlson, G., et al. (2005). Age-dependent neurofibrillary tangle formation, neuron loss, and memory impairment in a mouse model of human tauopathy (P301L). J Neurosci 25, 10637–10647.

Regehr, W.G. (2012). Short-term presynaptic plasticity. Cold Spring Harb Perspect Biol 4, a005702.

Rocher, A.B., Crimins, J.L., Amatrudo, J.M., Kinson, M.S., Todd-Brown, M.A., Lewis, J., and Luebke, J.I. (2010). Structural and functional changes in tau mutant mice neurons are not linked to the presence of NFTs. Exp Neurol 223, 385–393.

Rubin, D.B., Van Hooser, S.D., and Miller, K.D. (2015). The stabilized supralinear network: a unifying circuit motif underlying multi-input integration in sensory cortex. Neuron 85, 402–417.

Sanchez-Vives, M.V., Nowak, L.G., and McCormick, D.A. (2000). Cellular mechanisms of long-lasting adaptation in visual cortical neurons in vitro. J Neurosci 20, 4286–4299.

Santacruz, K., Lewis, J., Spires, T., Paulson, J., Kotilinek, L., Ingelsson, M., Guimaraes, A., DeTure, M., Ramsden, M., McGowan, E., et al. (2005). Tau suppression in a neurodegenerative mouse model improves memory function. Science 309, 476–481.

Sara, S.J. (2009). The locus coeruleus and noradrenergic modulation of cognition. Nat Rev Neurosci 10, 211–223.

Schindelin, J., Arganda-Carreras, I., Frise, E., Kaynig, V., Longair, M., Pietzsch, T., Preibisch, S., Rueden, C., Saalfeld, S., Schmid, B., et al. (2012). Fiji: an open-source platform for biological-image analysis. Nat Methods 9, 676–682.

Scullion, S.E., Barker, G.R.I., Warburton, E.C., Randall, A.D., and Brown, J.T. (2019). Muscarinic Receptor-Dependent Long Term Depression in the Perirhinal Cortex and Recognition Memory are Impaired in the rTg4510 Mouse Model of Tauopathy. Neurochem Res 44, 617–626.

Solomon, S.G., and Kohn, A. (2014). Moving sensory adaptation beyond suppressive effects in single neurons. Curr Biol 24, R1012–1022.

Spires, T.L., Orne, J.D., SantaCruz, K., Pitstick, R., Carlson, G.A., Ashe, K.H., and Hyman, B.T. (2006). Region-specific dissociation of neuronal loss and neurofibrillary pathology in a mouse model of tauopathy. Am J Pathol 168, 1598–1607.

Tang-Wai, D.F., Graff-Radford, N.R., Boeve, B.F., Dickson, D.W., Parisi, J.E., Crook, R., Caselli, R.J., Knopman, D.S., and Petersen, R.C. (2004). Clinical, genetic, and neuropathologic characteristics of posterior cortical atrophy. Neurology 63, 1168–1174.

Teyler, T.J., Hamm, J.P., Clapp, W.C., Johnson, B.W., Corballis, M.C., and Kirk, I.J. (2005). Longterm potentiation of human visual evoked responses. Eur J Neurosci 21, 2045–2050.

Webster, M.A. (2015). Visual Adaptation. Annu Rev Vis Sci 1, 547–567.

Wes, P.D., Easton, A., Corradi, J., Barten, D.M., Devidze, N., DeCarr, L.B., Truong, A., He, A., Barrezueta, N.X., Polson, C., et al. (2014). Tau overexpression impacts a neuroinflammation gene expression network perturbed in Alzheimer’s disease. PLoS One 9, e106050.

William, C.M., Andermann, M.L., Goldey, G.J., Roumis, D.K., Reid, R.C., Shatz, C.J., Albers, M.W., Frosch, M.P., and Hyman, B.T. (2012). Synaptic plasticity defect following visual deprivation in Alzheimer’s disease model transgenic mice. J Neurosci 32, 8004–8011.

Witton, J., Staniaszek, L.E., Bartsch, U., Randall, A.D., Jones, M.W., and Brown, J.T. (2016). Disrupted hippocampal sharp-wave ripple-associated spike dynamics in a transgenic mouse model of dementia. J Physiol 594, 4615–4630.

Yue, M., Hanna, A., Wilson, J., Roder, H., and Janus, C. (2011). Sex difference in pathology and memory decline in rTg4510 mouse model of tauopathy. Neurobiol Aging 32, 590–603.

Zhao, X., Kotilinek, L.A., Smith, B., Hlynialuk, C., Zahs, K., Ramsden, M., Cleary, J., and Ashe, K.H. (2016). Caspase-2 cleavage of tau reversibly impairs memory. Nat Med 22, 1268–1276.

